# Topic modeling reveals thermally partitioned and taxonomically distinct microbial subcommunities across prokaryotes and phytoplankton in the Laurentian Great Lakes

**DOI:** 10.64898/2026.06.06.730626

**Authors:** María D. Hernández Limón, Claire Donnat, Freddy Bunbury, Maureen L. Coleman

## Abstract

Identifying discrete microbial assemblages and their environmental drivers across multiple biological fractions simultaneously remains a central challenge in aquatic microbial ecology. We applied an integrated analytical pipeline built around Latent Dirichlet Allocation (LDA) to an eight-year 16S rRNA amplicon time series from the Laurentian Great Lakes, spanning four size-fractionated biological blocks — free-living prokaryotes, particle-associated prokaryotes, and small and large chloroplast-containing eukaryotes. LDA resolved ecologically coherent subcommunities whose taxonomic identity was consistently defined at the order and class level, with fingerprint taxa confirmed by discriminant analysis. Shannon entropy differences between blocks reflected fundamental differences in dispersal capacity and environmental filtering — free-living prokaryotes and large eukaryotes showed higher mixing than particle-associated prokaryotes and small eukaryotes. Temperature dominated environmental structuring across all blocks, assessed through Limma and random forest with SHAP, with secondary drivers differing by size fraction. Group Compositional Analysis jointly integrating all four blocks revealed that thermal stratification and lake chemistry organize microbial communities coherently across all size fractions simultaneously. Warm stratified and cold inversely-stratified waters harbored largely non-overlapping assemblages across all four blocks, with cold water specialists — including chemolithotrophic deep-branching lineages and silica-dependent diatoms — having no warm water equivalents.

**Importance:** The Laurentian Great Lakes are among the fastest warming lakes in the world, yet the microbial communities that drive their biogeochemical cycles remain poorly characterized across size fractions and thermal habitats. Using eight years of samples spanning all five Great Lakes, we show that free-living bacteria, particle-associated bacteria, small phytoplankton, and large phytoplankton all respond coherently to the same master environmental gradients — thermal stratification and lake chemistry — despite fundamental differences in organism size, trophic role, and sequencing protocol. Warm stratified and cold water habitats support distinct microbial communities, and the cold water specialists identified here have no warm water equivalents. As the Great Lakes warm and cold water habitats shrink, the microbial communities that depend on them will not simply become less abundant they will be replaced by fundamentally different assemblages.

## Introduction

Microbial communities exhibit biogeographical patterns shaped by environmental conditions across diverse aquatic systems, including oceans and lakes (Follows et a;., 2007, Hanson et al., 2012; Martiny et al., 2006). Identifying the ecological units that underlie these patterns — groups of taxa that co-occur due to shared environmental responses (De Wit & Bouvier, 2006) or compatible functions — is a central challenge in microbial ecology. Such subcommunities (SCs) can be defined without assuming direct interactions among taxa; instead, members co-occur through spatial, temporal, or functional partitioning (Blei et al. 2003; Sankaran & Holmes 2019). Defining SCs provides insight into microbial population structure, niche partitioning, and how distinct environmental gradients independently shape community assembly (Louca et al., 2018; Faust & Raes, 2012).

The Laurentian Great Lakes offer an ideal system for investigating these patterns. Spanning a large geographic gradient with pronounced seasonal variation in temperature, mixing, and nutrient dynamics, these lakes are among the fastest warming in the world (O’Reilly et al., 2015, Woolway et al., 2021; Ozersky et al., 2021). As stratification intensifies and cold water habitats contract under warming, understanding which microbial communities are at risk and which will expand becomes increasingly urgent. The five lakes also differ markedly in trophic status, from the oligotrophic waters of Lake Superior to the productive basins of Lakes Erie and Ontario, creating overlapping thermal and chemical gradients that may independently structure microbial communities. Despite the foundational role of microbes in biogeochemical cycling and primary production (Falkowski et al., 2008; Newton et al., 2011), microbial community research in the Great Lakes remains limited relative to studies of higher trophic levels. Emerging work has identified strong biogeographical patterns, including lake- and depth-specific communities (Bramburger & Reavie, 2016; Paver et al., 2020; Podowski et al., 2022; Gale et al., 2023), but whether thermal and trophic gradients represent distinct axes of subcommunity assembly and how these signals are expressed across microbial size fractions remains unknown.

Characterizing microbial subcommunity assembly across environmental gradients presents compounding methodological challenges. Standard co-occurrence approaches force taxa into mutually exclusive groups and cannot easily integrate community structure across datasets with different compositions (Faust, 2021; Kishore et al., 2023). We address these limitations through an integrated analytical pipeline built around Latent Dirichlet Allocation (LDA), a probabilistic topic modeling approach that identifies SCs as distributions of co-occurring taxa across samples, allowing individual taxa to contribute to multiple SCs (Sankaran & Holmes, 2019). Topic number is optimized using resolution and coherence metrics, and topic entropy characterizes the degree of mixing within samples (Fukuyama et al., 2023) — whether communities are dominated by a single SC or represent mixtures of multiple co-occurring assemblages. Fingerprint taxa defining each SC are identified using discriminant analysis (GoM-DE) (Carbonetto et al., 2023), and environmental associations are assessed through Limma (Ritchie et al., 2015), random forest (Breiman (2001), and SHAP (Lundberg & Lee, 2017), which together rank driver importance and directionality across all topics.

Because the four biological blocks — free-living prokaryotes (FL), particle-associated prokaryotes (PA), small size fraction chloroplasts (SSF), and large size fraction chloroplasts (LSF) — cannot be directly compared across size fractions due to differences in sequencing protocols, LDA projects each into a common latent space (Sankaran & Holmes 2019). Generalized canonical analysis (GCA) (Tenenhaus & Tenenhaus 2014); then jointly integrates all four topic distributions to reveal shared axes of cross-fraction covariation. Although LDA has been applied to soil and host-associated microbiomes (Kim et al., 2023; Pappalardo et al., 2024), this pipeline has not been previously applied to pelagic microbial communities. Using an eight-year 16S rRNA amplicon time series of the Laurentian Great Lakes (Supp. Fig. 1), we show that microbial subcommunities recover known ecological structure — including thermal stratification and lake identity as distinct axes of cross-fraction covariation — and reveal the taxonomic identities of the communities associated with each gradient. The warm stratified epilimnion and cold inversely-stratified habitats that define opposite ends of the dominant community axis harbor largely non-overlapping assemblages with no warm water equivalents for cold water specialists, a finding with direct implications for understanding how Great Lakes microbial communities will respond as stratification intensifies and cold water habitats contract under continued warming (O’Reilly et al., 2015; Woolway et al., 2021).

**Fig 1.**
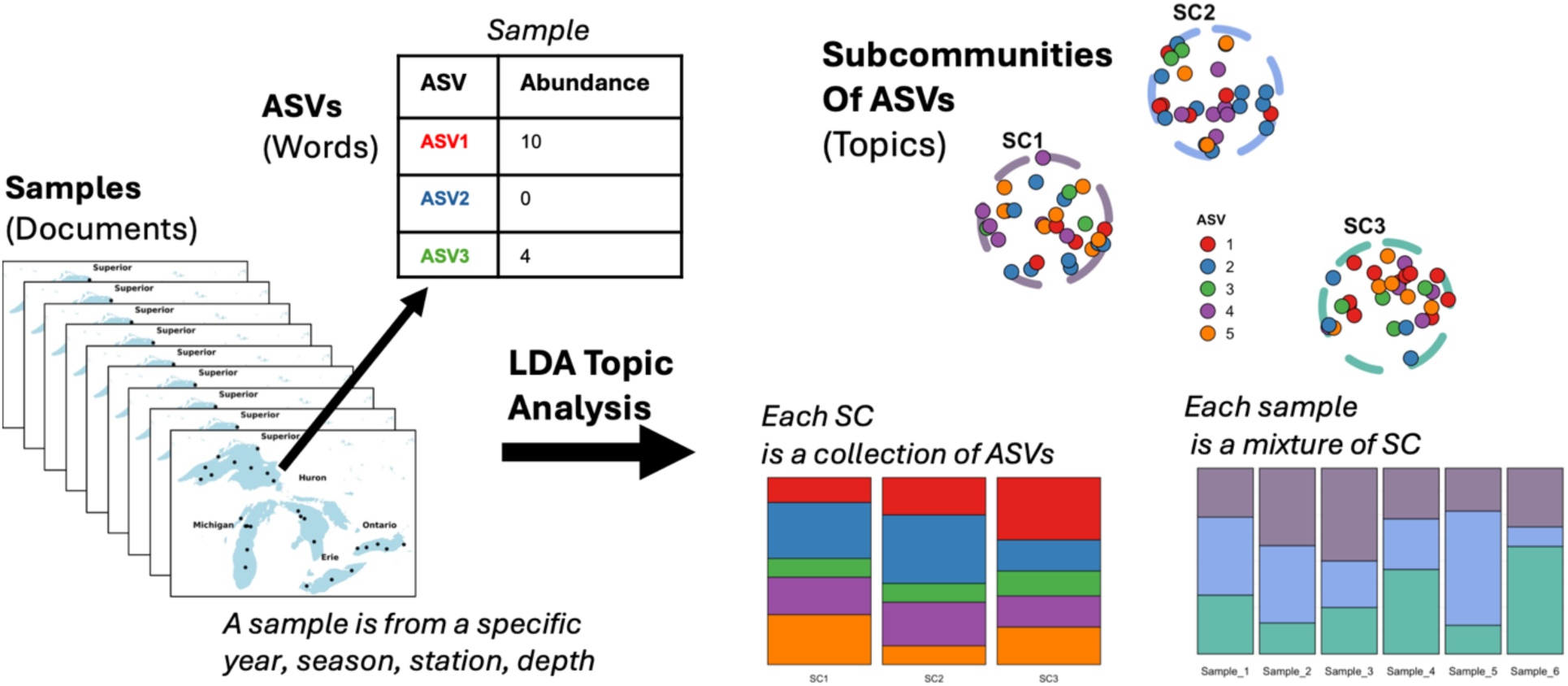
Conceptual diagram illustrating the application of Latent Dirichlet Allocation (LDA) to microbial community data from the Great Lakes. In this framework, each water sample is treated like a document, composed of words (taxa or ASVs) with varying abundances. LDA models the distribution of taxa across samples by identifying topics—interpreted here as microbial subcommunities (SC). Each subcommunity is a probabilistic mixture of taxa, and each sample is modeled as a mixture of these subcommunities or topics. This approach allows for dimensionality reduction and facilitates ecological interpretation of microbial community structure.

## Results and Discussion

### Subcommunities Represent Coherent Assemblages Across Space and Time

We applied Latent Dirichlet Allocation (LDA) separately to four size-fractionated datasets — free-living prokaryotes (FL, 0.2–1.6 µm), particle-associated prokaryotes (PA, >1.6 µm), and chloroplast-containing phytoplankton in small (SSF, 0.2–1.6 µm) and large (LSF, >1.6 µm) size fractions. The number of subcommunities (SCs) was determined using dataset-specific diagnostic criteria (see Methods, Supp Fig. 2) resulting in 10 SCs for FL and PA, 9 for SSF, and 11 for LSF. In this framework, each SC represents a reproducible set of co-occurring taxa with compatible life strategies or overlapping niches (Faust & Raes, 2012; Louca et al., 2018), and each sample is modeled as a mixture of SCs in varying proportions (Fig. 1, Fig. 2A). For example, a spring sample from western Lake Erie might reflect contributions from a ubiquitous freshwater SC, a riverine input SC, a spring bloom-associated SC, and a resuspended sediment SC — each weighted differently depending on local conditions and interannual variability. In the Laurentian Great Lakes, where hydrological connectivity facilitates widespread dispersal (Sterner, 2021), this mixture-based framework captures how entire assemblages shift along environmental gradients rather than simply tracking the presence or absence of individual taxa.

**Fig. 2.**
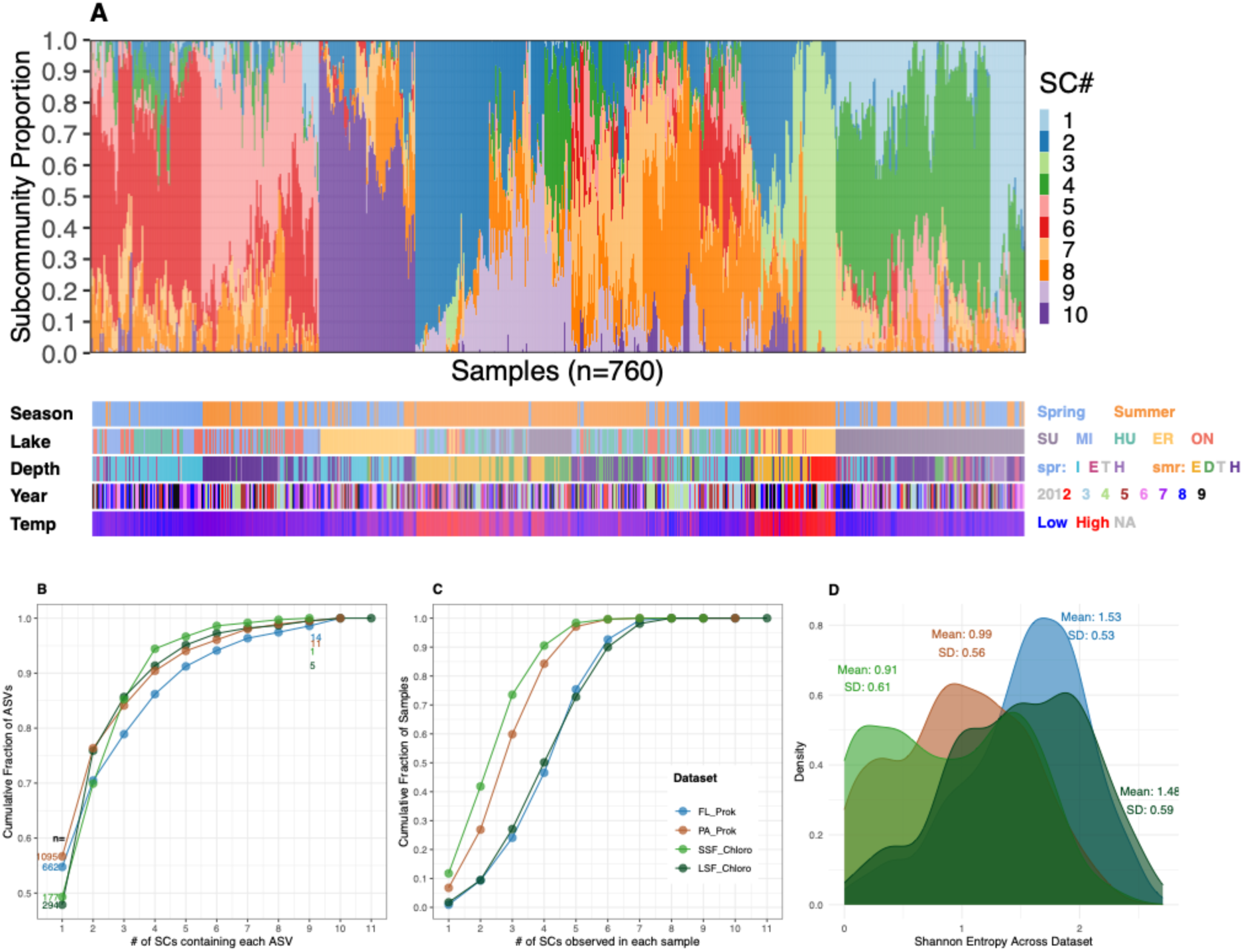
Subcommunity organization across biological blocks. (A) Sample-by-subcommunity (SC) composition for free-living prokaryotes (FL, n=760), with samples ordered by Ward clustering and colored by SC identity. Metadata tracks below show season, lake, depth, year, and temperature. (B) Cumulative distribution of AS Vs by the number of SCs in which they were detected (p>0.0001) across all four size fractions; n values indicate the number of ASVs found in exactly 1 or 9 SCs (the highest number of SCs all datasets have). (C) Cumulative distribution of samples by the number of SCs detected per sample (p>0.01). (D) Shannon entropy of SC composition per sample for each dataset.

### Subcommunity Organization Differs Systematically Across Biological Blocks

LDA resolved biologically meaningful subcommunities across all four size-fractionated datasets (Supplementary Fig. 3-4). As illustrated for free-living prokaryotes (Fig. 2A), SC composition tracked coherently with season, lake identity, depth, and temperature — structure that emerged from taxon co-occurrence patterns alone without any environmental information supplied to the model. This coherence with metadata confirms that the recovered SCs represent ecologically structured assemblages rather than statistical artifacts. To characterize how SCs are organized across biological blocks, we examined both taxon-level SC specificity and sample-level SC mixing. Most ASVs were associated with a single SC regardless of fraction (Fig. 2B), indicating that taxon-level specialization is a consistent feature across all blocks. However, sample-level mixing differed markedly across biological blocks — Shannon entropy of SC composition was higher in free-living prokaryotes (FL; mean ± SD: 1.53 ± 0.53) and large size fraction chloroplasts (LSF; 1.48 ± 0.59) than in particle-associated prokaryotes (PA; 0.99 ± 0.56) and small size fraction chloroplasts (SSF; 0.91 ± 0.61), consistent with the SC distribution across samples (Fig. 2C, D). These contrasting entropy patterns were statistically significant: FL and PA differed markedly in their SC composition distributions (Kolmogorov-Smirnov test: D = 0.41, p < 0.001), as did SSF and LSF (D = 0.34, p < 0.001), confirming that the differences in sample-level mixing between free-living and particle-associated fractions, and between small and large eukaryotes, are not attributable to chance.

These contrasting entropy patterns likely reflect fundamental differences in microbial lifestyle and niche breadth. This free-living versus particle-associated divergence in community organization is consistent with global patterns observed across marine and freshwater systems, where particle-associated communities consistently show greater environmental specificity and lower overlap with free-living assemblages (Mestre et al., 2018; Salazar et al., 2015; Rösel et al., 2012; Fujimoto et al., 2016). FL communities are characterized by greater dispersal potential and ecological generalism, producing more fluid, well-mixed assemblages shaped by environmental variability (Dang & Lovell, 2015). LSF communities similarly show broad taxonomic mixing, consistent with size-structured ecological breadth in larger phytoplankton (Litchman & Klausmeier, 2008; Hillebrand et al., 2022; Irwin et al., 2006). In contrast, PA and SSF communities tend to be more compositionally discrete and environmentally constrained — particle microenvironments impose steep oxygen gradients, complex substrates, and high surface-area heterogeneity that favor taxa with specialized metabolic capabilities or surface-adapted lifestyles (Azam & Long, 2001; Grossart, 2010; Fontanez et al., 2015), while small phytoplankton often exhibit strong environmental specialization tied to nutrient and light availability (Callieri et al., 2002; Callieri & Stockner, 2002; Worden et al., 2004). Together these patterns highlight how size fraction, particle association, and dispersal capacity jointly shape subcommunity organization across the Great Lakes, independently of the number of SCs recovered by LDA.

### Subcommunities Are Taxonomically Distinct and Environmentally Structured Across All Biological Blocks

To characterize the taxonomic composition of each subcommunity, we applied GoM-DE discriminant analysis (Carbonetto et al., 2023) to identify ASVs differentially enriched in each SC across all four biological blocks. GoM-DE analysis identified differentially enriched ASVs across all four biological blocks, yielding 2,688 fingerprint ASVs in FL (Fig. 3B), 2,044 in PA (Supp. Fig. 5), 193 in SSF (Supp. Fig. 6), and 407 in LSF (Supp. Fig. 7). Consistent with the ASV distribution shown in Fig. 2B, the vast majority of fingerprint ASVs were enriched in exactly one SC across all blocks (Fig. 3B, Supp. Fig. 5-7) — confirming that LDA-recovered SCs represent biologically distinct assemblages rather than statistical groupings. The signal strength of this enrichment varied across SCs, with some showing markedly high log2FC values indicating strong environmental selection for a specific set of co-occurring taxa (Fig. 3C), while others showed weaker differentiation consistent with broader environmental tolerance or transitional community states (Supp. Fig. 5–7).

**Fig. 3.**
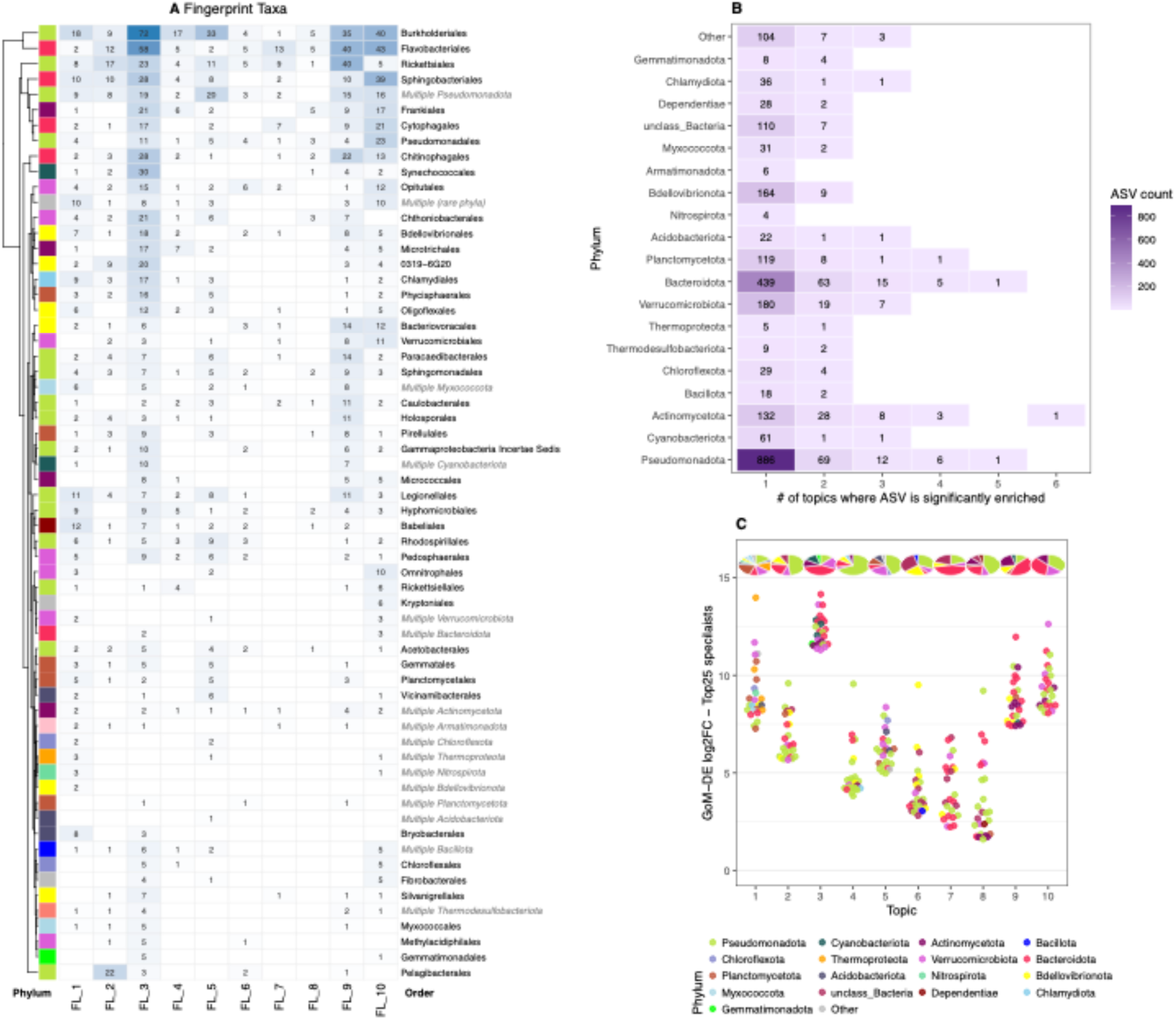
GoM-DE fingerprint taxa and subcommunity specificity for free-living prokaryotes. (A) Fingerprint taxa heatmap showing the number of differentially enriched ASVs per taxonomic order across all 10 FL subcommunities (SCs), restricted to ASVs enriched in exactly one SC (see panel B). Rows are taxonomic orders clustered by phylum identity (dendrogram); row label colors distinguish named orders (black) from phylum-level collapsed entries (grey italic, e.g., Multiple Pseudomonadota). Values indicate ASV counts per SC. Row/phylum colors correspond to the legend in panel C. Full GoM-DE results for PA, SSF, and LSF are shown in Supplementary Fig. 6-9. (B) Distribution of all marker ASVs (n=2,688) by the number of SCs in which each ASV was significantly enriched. (C) log2 fold-change of the top 25 specialist ASVs per SC. Points are colored by phylum; pie charts above each SC show the phylum composition of all specialist ASVs for that topic.

**Fig. 4.**
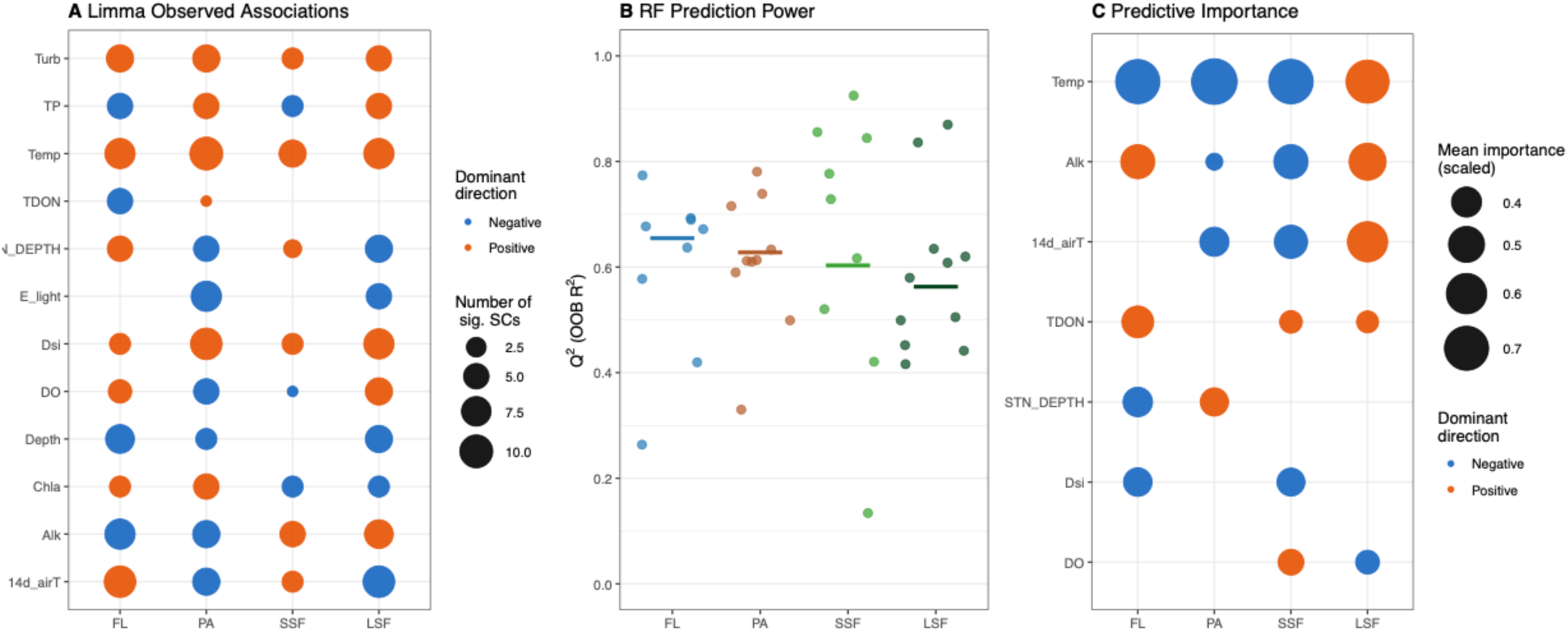
Environmental structuring of subcommunities across biological blocks. (A) Limma observed associations between environmental variables and SC composition across all four biological blocks. Circle size indicates the number of SCs with a statistically significant association (BH-adjusted p < 0.05); color indicates dominant direction of the t-statistic across significant SCs (orange = positive, blue = negative). Full per-SC Limma results for each block are shown in Supplementary Fig. 6-9C. (B) RF prediction power (Q^2^; OOB R^2^) for each SC across all four biological blocks. Each point represents one SC; horizontal bars indicate block means. (C) Cross-block summary of RF+SHAP predictive importance. Circle size reflects mean scaled SHAP importance across SCs within each block; color indicates dominant direction derived from SHAP value correlations (orange = positive, blue = negative) where at least 60% of SCs within a block agree. Full RF and SHAP results per block are shown in Supplementary Fig. 6-9D.

**Fig. 5.**
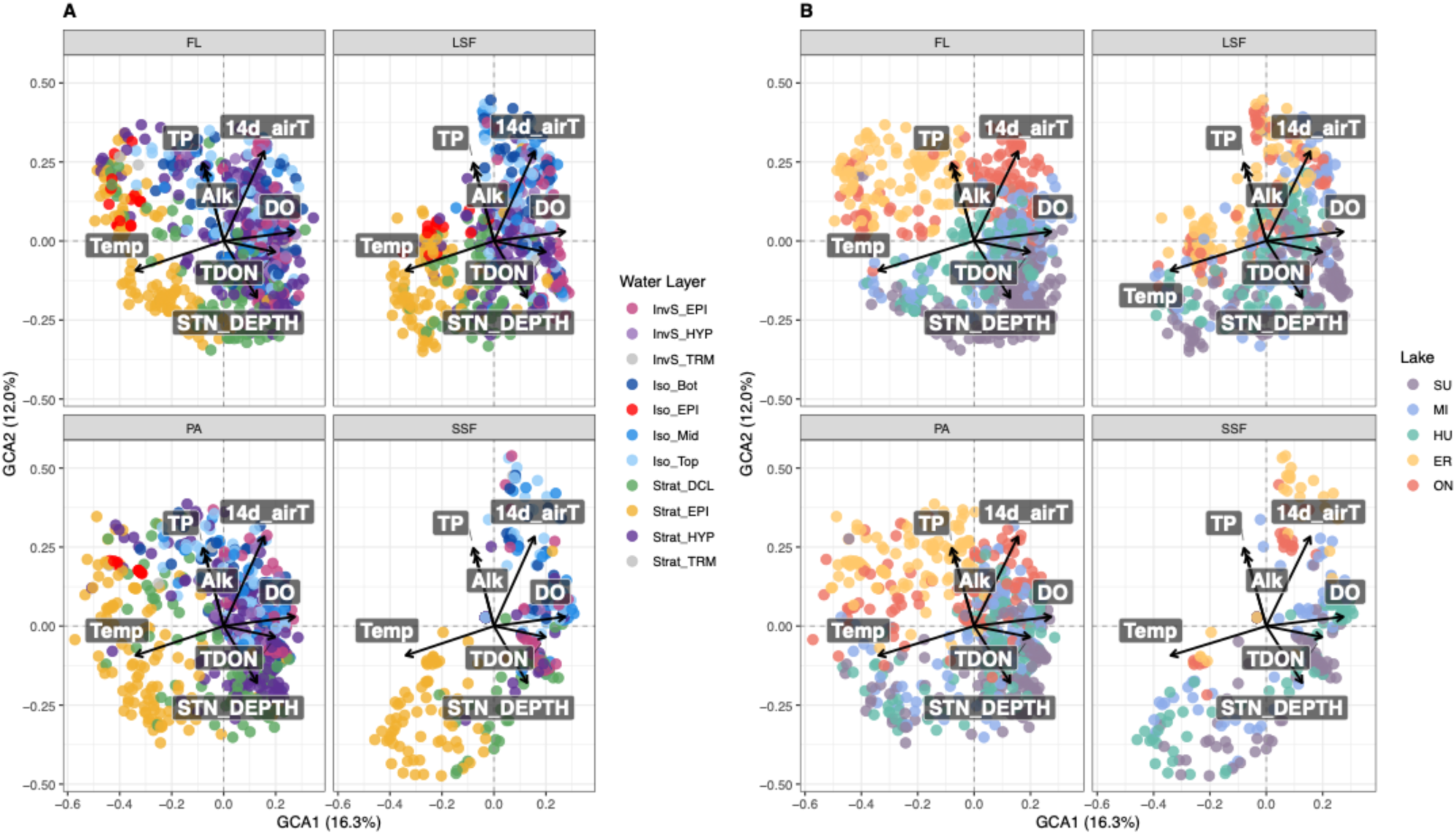
Group Compositional Analysis (GCA) biplots across all four biological blocks. Sample scores on GCA1 and GCA2 are shown for free-living prokaryotes (FL), particle-associated prokaryotes (PA), small size fraction chloroplasts (SSF), and large size fraction chloroplasts (LSF). Environmental vectors show the direction and relative magnitude of environmental associations with GCA axes; only the longest 80% of vectors are shown for clarity. (A) Samples colored by water layer. (B) Samples colored by lake. Block contributions and environmental loadings for all five axes are shown in Supplementary Fig. 15.

**Fig. 6.**
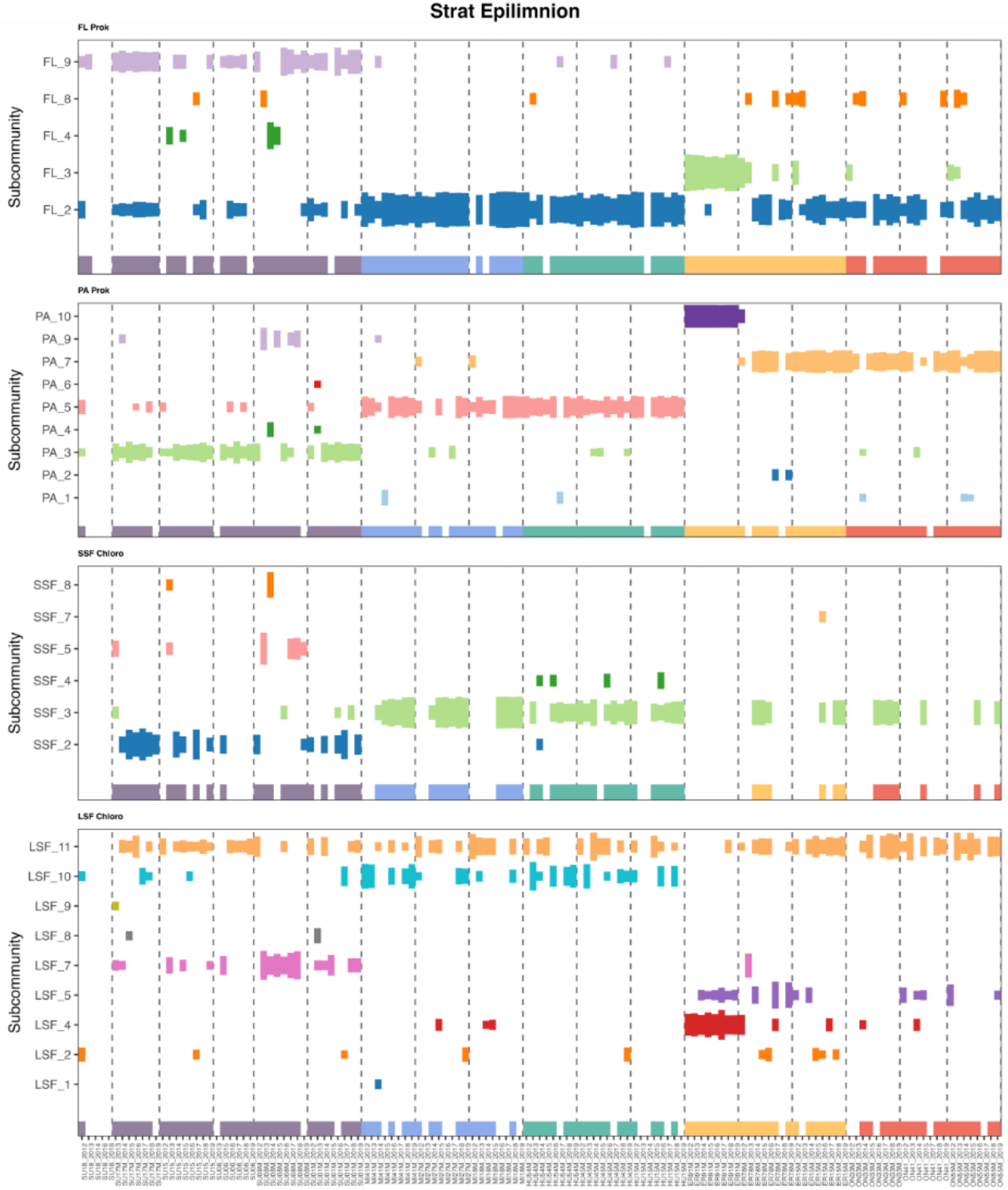
Subcommunity composition of warm stratified epilimnion waters. SC proportional abundance across samples for all four biological blocks, ordered from western Lake Superior to eastern Lake Ontario stations with years 2012-2019 nested within each station (dashed lines). Only SCs contributing more than three times the uniform expectation (3/K, where K is the number of SCs) are shown.

**Fig. 7.**
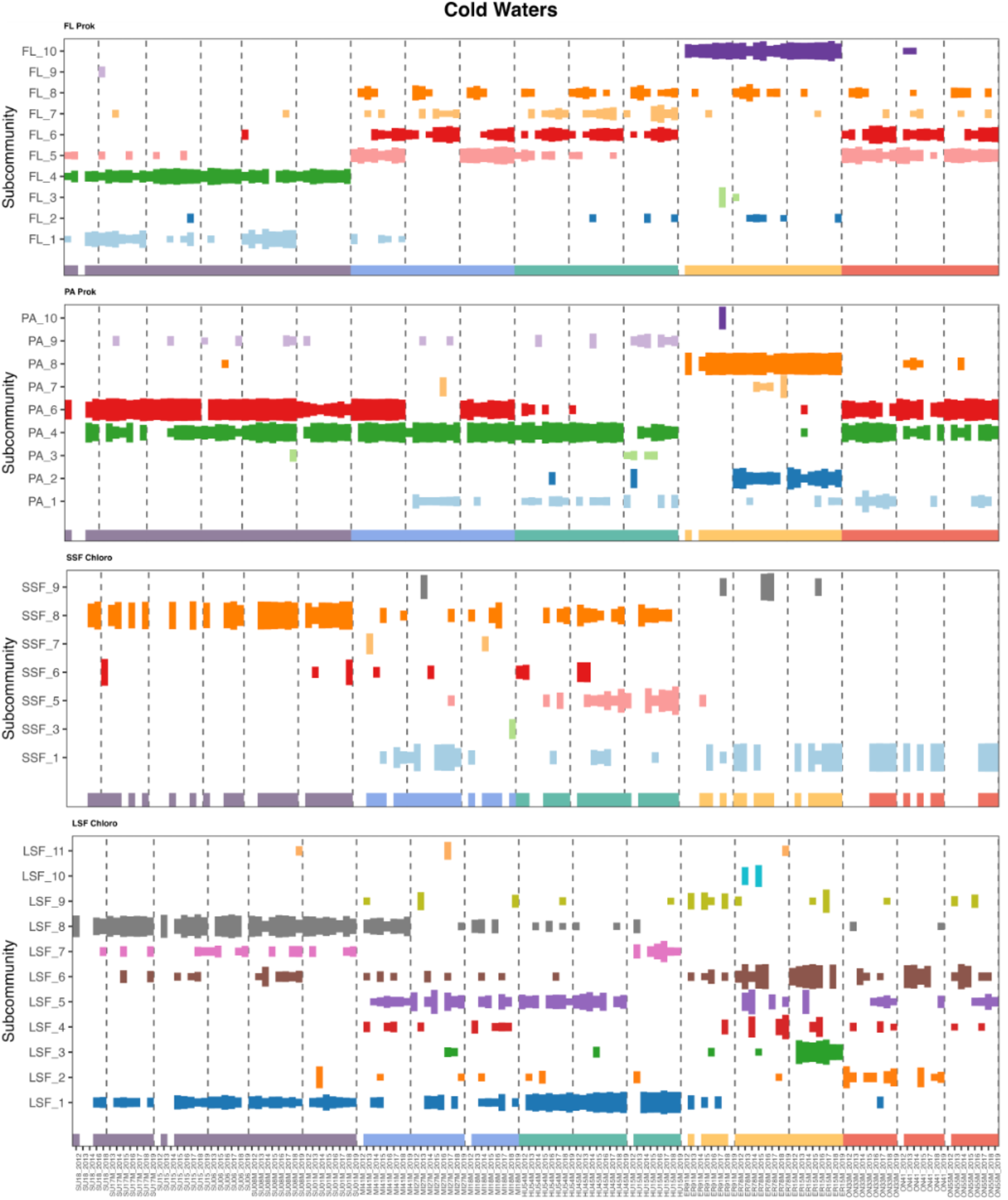
Subcommunity composition of cold water habitats. SC proportional abundance across samples for all four biological blocks, ordered from western Lake Superior to eastern Lake Ontario stations with years 2012-2019 nested within each station (dashed lines). Cold water samples include stratified hypolimnion, spring isothermal, and spring inversely-stratified conditions. Only SCs contributing more than three times the uniform expectation (3/K, where K is the number of SCs) are shown.

While members of the same phyla appeared across multiple SCs in all blocks, SC identity was consistently defined at finer taxonomic resolution — specific orders and classes differed even when the same phylum contributed fingerprint ASVs to several SCs (Fig. 3A, Supp. Fig. 8-11A).

Pseudomonadota contributed the largest number of fingerprint ASVs in both FL (886) and PA (677), but despite this numerical dominance the majority were enriched in a single SC, indicating that while Pseudomonadota are taxonomically diverse across the community, individual ASVs maintain highly specific SC associations (Fig. 3, Supp. Fig. 5). Bacteroidota contributed fingerprint ASVs across nearly all SCs in both FL and PA and showed the highest proportion of multi-SC ASVs among all phyla (Fig. 3B, Supp. Fig. 5), consistent with their known ecological generalism as dissolved organic matter degraders (Cottrell & Kirchman, 2000; Teeling et al., 2012). Despite substantial overlap in dominant orders between FL and PA — particularly Burkholderiales, Flavobacteriales, and Sphingobacteriales — PA fingerprint taxa showed greater representation of particle-associated lineages including Planctomycetota and Rhodobacterales, while FL retained free-living lineages such as Pelagibacterales (Brown et al., 2012) and Synechococcales (Yeh & Fuhrman, 2022), consistent with the selective pressure of particle microenvironments on community composition (Mestre et al., 2018; Grossart, 2010; Salazar et al., 2015). Among eukaryotic fractions, Chrysophyceae emerged as the dominant fingerprint class in both SSF and LSF (Supp. Fig. 6-7, 10-11), suggesting size-structured populations of the same functional group maintaining SC-specific associations across both size fractions. LSF additionally captured larger taxa absent from SSF — including Chlamydomonadales, Euglenida, and Dictyochophyceae — reflecting the size-dependent partitioning of the phytoplankton community across fractions (Reynolds, 2006; Hillebrand et al., 2022; Irwin et al., 2006). Together these patterns demonstrate that LDA recovers biologically meaningful subcommunity structure organized at the order and class level, even within phyla that span multiple SCs.

Observed associations between SC composition and environmental parameters were widespread and statistically significant across all four biological blocks (Fig. 4A, Supp. Fig. 8-11). Temperature showed the strongest and most consistent associations, with significant relationships detected across the greatest number of SCs in all four blocks — a pattern reinforced by RF+SHAP predictive importance (Fig. 4C), confirming temperature as the dominant driver of SC composition across the Great Lakes microbial system (Shade et al., 2007; Shade et al., 2012; Yannarell & Triplett, 2004). Alkalinity showed significant associations across all blocks but with opposing directions between prokaryotic and eukaryotic fractions, likely reflecting its role as a proxy for lake identity — particularly the distinctively low alkalinity of Lake Superior relative to the other Great Lakes — rather than a direct physiological effect on community composition (Sterner, 2021). Air temperature (14d_airT) emerged as an important secondary driver across PA, SSF, and LSF but was less prominent in FL, suggesting that larger and particle-associated communities track atmospheric thermal forcing more directly than free-living bacterioplankton. This may reflect the longer generation times and slower turnover of particle-associated and larger eukaryotic communities, which integrate thermal signals over longer timescales than the rapidly dividing free-living bacterioplankton (Grossart, 2010). Dissolved silica and turbidity showed block-specific associations consistent with their known roles in structuring diatom-associated communities and productive lake environments respectively (Bramburger & Reavie, 2016; Reavie et al., 2025).

RF models predicted SC composition with high accuracy across all blocks (Fig. 4B), with mean Q² values of 0.65 (FL), 0.63 (PA), 0.60 (SSF), and 0.56 (LSF). The decline in predictability from FL to LSF is consistent with free-living prokaryotes being most strongly filtered by measured environmental gradients, while larger eukaryotes may be subject to additional biotic interactions or stochastic processes not captured by the 12-variables selected. Together the Limma and RF+SHAP results confirm that the subcommunities recovered by LDA are not only taxonomically distinct but environmentally coherent — the high RF predictability across all blocks suggesting that SC composition is deterministically structured by the measured environmental gradients rather than stochastically assembled, consistent with environmental filtering as the dominant force in Great Lakes microbial community assembly (Hanson et al., 2012).

### Stratification and Lake Identity Dominate Cross-Block Microbial Organization with Block-Specific Secondary Axes

Having characterized subcommunity organization and environmental drivers within each biological block independently, we next asked whether coherent patterns emerge when all four blocks are considered simultaneously. The four datasets derive from two size fractions — free-living (FL + SSF) and particle-associated (PA + LSF) — with prokaryotic and chloroplast communities sequenced together within each fraction. While FL and SSF are directly comparable, as are PA and LSF, cross-fraction comparisons are complicated by both the fundamental ecological differences between size fractions and differences in sequencing protocols — even where community composition distributions are broadly similar (Hernández Limón et al., *in review*). Direct comparisons across size fractions are therefore not appropriate. LDA addresses this limitation by projecting each dataset into a common latent space of subcommunity proportions, enabling Group Compositional Analysis (GCA) to jointly integrate all four topic distributions and identify shared axes of variation across the full microbial system.

GCA jointly integrated the subcommunity distributions across all four biological blocks, identifying shared axes of variation that transcend individual size fractions. The first two axes explained 16.3% (GCA1) and 12.0% (GCA2) of total shared variance respectively, with all four biological blocks contributing meaningfully to both axes (Supp. Fig. 14A). The combined 28.3% of shared variance explained by GCA1 and GCA2 reflects the inherent complexity of integrating four biologically distinct datasets — the fact that two interpretable axes emerge consistently across all blocks is itself a strong result.

GCA1 captured water column stratification — consistent with the dominant axis recovered by beta diversity analyses in the Laurentian Great Lakes and other large lake systems (Hernández Limón et al., *in review*). Across all four blocks, warm stratified epilimnion samples consistently occupied positive GCA1 scores while cold inversely stratified and isothermal samples occupied negative scores (Fig. 5A), with temperature loading strongly negative and dissolved oxygen and air temperature loading positively (Supp. Fig. 12B). What is notable here is not the stratification signal itself — well established from beta diversity approaches including independent ordinations of the FL and PA fractions (Hernández Limón et al., *in review*) — but that this signal emerges coherently across all four biological blocks simultaneously, derived purely from taxon co-occurrence patterns without any environmental information supplied to the model. That this stratification signal emerges from co-occurrence patterns alone validates the biological meaningfulness of the LDA framework and demonstrates that microbial community composition encodes environmental information recoverable through unsupervised topic modeling. This cross-block coherence could not have been demonstrated by analyzing each size fraction independently.

GCA2 (12.0%) captured lake identity, distinguishing the oligotrophic upper lakes (Superior, Michigan, Huron) from the more productive lower lakes (Erie, Ontario) along a gradient of total phosphorus and alkalinity (Fig. 5B, Supp. Fig. 12B). Alkalinity in this system functions primarily as a proxy for lake identity rather than a direct physiological driver, reflecting the well-documented differences in nutrient loading, water residence time, and carbonate chemistry between the upper and lower Laurentian Great Lakes (Sterner et al., 2021). The consistency of this upper vs lower lake separation across all four biological blocks suggests that lake trophic identity is a stronger organizer of microbial community composition than previously recognized from single-fraction studies.

GCA3 (8.3%) and GCA4 (5.8%) explained progressively less shared variance and captured more block-specific signals. GCA3 was dominated by SSF and associated with depth, station depth, and air temperature loadings, consistent with the strong vertical structuring of small phytoplankton communities — particularly their association with the deep chlorophyll layer (Bramburger & Reavie, 2016) — that is not shared as strongly with the prokaryotic fractions. GCA4 appeared to capture a PA-specific signal associated with turbidity and alkalinity, consistent with particle-associated communities in productive and turbid lake environments (Grossart, 2010; Mestre et al., 2018). GCA5 (4.9%) similarly captured a block-specific signal dominated by LSF and loading strongly on chlorophyll and estimated light availability (Supp. Fig. 12), consistent with the light-dependent growth strategies of larger phytoplankton (Litchman & Klausmeier, 2008; Irwin et al., 2006). The emergence of these block-specific axes is consistent with the differences in subcommunity organization observed in Fig. 2 — SSF’s strong depth structuring (GCA3) parallels its lower entropy and more discrete SC composition, while LSF’s light-driven axis (GCA5) reflects the greater sensitivity of larger phytoplankton to vertical light gradients. Together GCA3, GCA4, and GCA5 suggest that while stratification and lake chemistry organize all four biological blocks coherently along the first two axes, each biological block retains unique axes of variation reflecting its distinct ecological niche.

### Warm and Cold Waters Harbor Non-Overlapping Microbial Assemblages Across All Biological Blocks

GCA1 identified thermal stratification as the dominant axis of cross-block microbial organization, separating warm stratified epilimnion from cold inversely stratified and isothermal waters. To characterize the biological communities associated with each end of this gradient, we examined SC composition in stratified epilimnion (Strat_EPI) and cold water samples (stratified hypolimnion, spring isothermal, and spring inversely-stratified) separately across all four biological blocks (Figs. 6–7). Samples are ordered from the westernmost Lake Superior stations to the easternmost Lake Ontario stations, with years 2012–2019 nested within each station.

The most striking observation across both thermal regimes is that warm and cold waters harbor largely non-overlapping sets of subcommunities across all four biological blocks. Very few SCs appeared prominently in both thermal regimes, and where they did — such as FL_4 and FL_8 — their proportional contribution was markedly stronger in cold waters, indicating directional thermal preference rather than true thermal neutrality. This thermal partitioning was consistent across free-living prokaryotes, particle-associated prokaryotes, small and large eukaryotes alike, confirming that GCA1 captures genuine biological turnover between thermal habitats rather than a statistical artifact of the integrated analysis. The near-complete turnover of SC membership between warm and cold thermal regimes implies that warming-driven expansion of stratified epilimnion conditions (O’Reilly et al., 2015; Woolway et al., 2021) would not simply shift the relative abundance of existing communities but would replace cold water assemblages with taxonomically distinct warm water ones — a qualitative rather than quantitative change in community composition. Within each thermal regime, additional lake-specific structure was evident — reflecting the GCA2 lake chemistry signal — with distinct SCs characterizing Superior, Michigan/Huron, and Erie/Ontario in both warm and cold conditions. The persistence of lake-specific structure within each thermal regime confirms that thermal stratification and lake chemistry operate as independent axes of community assembly, consistent with the distinct GCA1 and GCA2 axes identified by the integrated analysis.

### Warm Stratified Surface Waters Support Lake-Structured Epilimnetic Assemblages Across All Biological Blocks

Stratified epilimnion samples revealed a consistent set of co-occurring subcommunities across all four biological blocks (Fig. 6). Warm surface waters were characterized by Pelagibacterales and Rickettsiales in the free-living fraction (FL_2 — the single largest SC in the FL dataset), Flavobacteriales, Burkholderiales, and Chitinophagales in the particle-associated fraction, and Chrysophyceae in both small and large eukaryotic fractions (SSF_2, SSF_3, LSF_11). While FL and LSF supported a cosmopolitan background SC alongside lake-specific SCs, PA warm SCs were lake-specific throughout yet taxonomically convergent at the order level. Lake identity was encoded in the finer taxonomic composition across blocks: in the free-living fraction, the lower lake warm SC (FL_3) was strongly dominated by Burkholderiales and included Synechococcales — suggesting a Cyanobacteria-associated heterotroph signal consistent with more productive lower lake conditions — while the upper lake warm SC (FL_9) showed a more balanced heterotroph assemblage with prominent Bacteriovoracales and Verrucomicrobiales. In the particle-associated fraction, Synechococcales characterized Superior warm waters (PA_3), Flavobacteriales, Burkholderiales, and Chitinophagales dominated Michigan and Huron (PA_5), Cyanobacteriales and Synechococcales were prominent in Erie and Ontario (PA_7), and elevated Chitinophagales alongside Synechococcales characterized the deep Erie station (PA_10). Among eukaryotic fractions, the warm SSF signal was sparse and Chrysophyceae-dominated across both SU (SSF_2, with only Chrysophyceae_X and Mamiellophyceae) and non-SU lakes (SSF_3), consistent with the strong environmental filtering of small phytoplankton in warm surface waters (Callieri et al., 2002; Worden et al., 2004). LSF warm SCs showed greater taxonomic breadth — LSF_11 combined Chrysophyceae_X with green algae including Sphaeropleales and Chlamydomonadales across all lakes, while lower lake warm waters additionally showed a Cryptophyceae signal (LSF_5) absent from the upper lakes.

The taxonomic composition of warm surface subcommunities reflects well-established ecological strategies for life in stratified, sunlit surface waters. The dominance of Pelagibacterales and Rickettsiales in FL_2 is consistent with the success of streamlined, oligotrophic bacterioplankton in stable surface environments — these lineages are characterized by minimal nutrient requirements and broad environmental tolerance, enabling their cosmopolitan distribution across all five lakes (Brown et al., 2012). The co-occurrence of Burkholderiales and Synechococcales in the lower lake warm SC (FL_3) suggests active phytoplankton-heterotroph coupling — Synechococcus-driven primary production fueling copiotrophic Burkholderiales through dissolved organic matter release, a dynamic well-documented in productive freshwater and marine systems (Teeling et al., 2012; Needham & Fuhrman, 2016). In contrast, the prominence of Bacteriovoracales in the upper lake warm SC (FL_9) points to active top-down predatory control of bacterioplankton, more pronounced in oligotrophic systems where prey diversity supports specialist predators (Davidov & Jurkevitch, 2004; Pernthaler, 2005). The association of Cyanobacteriales with particle-associated warm SCs in Erie and Ontario (PA_7) is consistent with bloom-associated particle colonization in productive lower lake waters (Vanderploeg et al., 2010; Yeh & Fuhrman, 2022). Among eukaryotes, the dominance of Chrysophyceae across both SSF and LSF warm SCs reflects the competitive advantage of mixotrophic flagellates in stratified, nutrient-limited surface waters — able to supplement photosynthesis with phagotrophy when inorganic nutrients are limiting (Bock et al., 2022; Litchman & Klausmeier, 2008). The additional presence of green algae including Sphaeropleales and Chlamydomonadales in LSF_11 is consistent with their known preference for warm, stable, stratified conditions across temperate lakes (Reynolds, 2006; Litchman et al., 2009). Together the warm surface assemblages across all four biological blocks — spanning streamlined oligotrophs, copiotrophic heterotrophs, predatory bacteria, mixotrophic flagellates, and green algae — represent the communities positioned to expand as stratification intensifies and the warm epilimnion deepens and persists longer under warming (O’Reilly et al., 2015; Woolway et al., 2021).

### Cold Water Habitats Harbor Taxonomically Distinct Assemblages with No Warm Water Equivalents

While warm surface assemblages represent a seasonally structured overlay, cold water habitats — stratified hypolimnion, spring isothermal, and spring inversely-stratified samples — harbor the dominant microbial signal across PA, SSF, and LSF. This contrasts with the free-living prokaryote fraction where the cosmopolitan warm surface SC (FL_2) dominates the dataset despite equivalent sampling across depths and seasons — suggesting that the cold water dominance in particle-associated and eukaryotic fractions reflects genuine biological differences in community biomass and persistence rather than a sampling artifact. This pattern suggests that particle-associated and eukaryotic communities are more tightly coupled to the physical persistence of cold water conditions — their biomass and SC dominance reflecting the greater volumetric extent of cold water habitats in these deep, thermally stratified lakes (Sterner, 2021) — rather than the seasonal intensity of warm surface conditions. Cold water SCs were markedly more lake-specific than their warm water counterparts across all four biological blocks (Fig. 7), with FL_2 and FL_3 essentially absent from cold waters, confirming the thermal partitioning observed across the full dataset.

Superior and upper lake cold waters harbored the most taxonomically distinctive assemblages across all blocks. In the free-living fraction, Superior and Michigan cold SCs (FL_1, FL_4) were characterized by rare deep-branching lineages — Planctomycetota, Chloroflexota, Nitrospirota, and Thermoproteota alongside Burkholderiales and Actinomycetota — chemolithotrophic and oligotrophic-adapted taxa essentially absent from all other SCs, consistent with Superior’s cold, deep, silica-rich waters selecting for highly specialized metabolic guilds (Karner et al., 2001; Pollet et al., 2014; Fujimoto et al., 2016; Rösel et al., 2012). These cold-adapted lineages extended into the widely distributed upper lake cold SC (FL_5), which harbored Chloroflexota and Thermoproteota alongside Burkholderiales — suggesting thermal regime rather than trophic status determines the distribution of these lineages. In the particle-associated fraction, the widely distributed cold SC (PA_6) was dominated by Burkholderiales and Planctomycetota alongside Hyphomicrobiales and Sphingomonadales — consistent with complex organic matter degradation on particle surfaces in cold stratified waters (Pollet et al., 2014; Grossart, 2010). Among eukaryotes, upper lake cold waters were characterized by Chrysophyceae and diatoms across both size fractions — SSF_8 combining Chrysophyceae_X with Chlorophyceae, and LSF_8 showing the strongest cold diatom signal with Diatomeae_XX and Cryptophyceae —while

LSF_1 was dominated by flagellates including Chrysophyceae, Dictyochophyceae, and Phaeocystales — the latter known for forming mucilaginous colonies and contributing to carbon export (Walker & Trimborn, 2024).

Erie cold waters presented a strikingly different biological portrait. The free-living cold SC (FL_10) was strongly copiotrophic — dominated by Flavobacteriales, Sphingobacteriales, and Burkholderiales alongside Nitrospirota — reflecting Erie’s more productive cold water environment with active nitrogen cycling (Podowski et al., 2022). Within the particle-associated fraction, Erie cold waters were further partitioned into two distinct niches — PA_2, associated with the stratified hypolimnion and positively correlated with total phosphorus and negatively with light, and PA_8, found predominantly in cold spring waters and positively associated with light — illustrating how within-lake environmental gradients structure communities at a finer scale than the broad GCA axes alone can capture. Despite this within-lake partitioning, both Erie cold PA SCs were dominated by Flavobacteriales, Burkholderiales, Chitinophagales, and Sphingobacteriales — a copiotrophic signal consistent across Erie’s cold water column. Among eukaryotes, cold Erie waters showed no distinctive SSF signal — consistent with the absence of a Superior-equivalent small phytoplankton cold community — while LSF cold SCs in the lower lakes were characterized by Cryptophyceae and Diatomeae_XX.

The taxonomic composition of cold water subcommunities across all four biological blocks reflects a fundamental contrast between the oligotrophic cold waters of the upper lakes and the productive cold waters of Erie. Superior and upper lake cold assemblages are distinguished by rare chemolithotrophic and deep-branching lineages — Chloroflexota, Thermoproteota, Nitrospirota, Planctomycetota — that are essentially absent from warm waters and from Erie cold waters, suggesting these taxa are cold-adapted specialists shaped by the unique physicochemical conditions of deep, silica-rich, oligotrophic environments. Erie cold assemblages in contrast are dominated by copiotrophic heterotrophs and active nitrogen cyclers, reflecting the influence of nutrient loading and organic matter availability on cold water community assembly even outside the warm stratified season. Among eukaryotes, the dominance of Chrysophyceae across cold SSF SCs confirms their broad thermal tolerance and mixotrophic flexibility (Bock et al., 2022; Callieri & Stockner, 2002), while the emergence of Diatomeae_XX in cold LSF reflects their silica-dependent, bloom-forming ecology during spring isothermal mixing (Reynolds, 2006; Bramburger & Reavie, 2016; Reavie et al., 2025; Barbiero et al., 2006). Together these patterns demonstrate that cold water habitats are not a single biological state but a diverse set of lake-specific assemblages — and that the organisms at risk as cold water habitats contract under warming are taxonomically irreplaceable, with no warm water equivalents.

## Conclusions

Together these results demonstrate that LDA-based subcommunity analysis recovers ecologically meaningful structure across four biologically distinct microbial size fractions simultaneously — identifying the same physicochemical gradients previously recovered through arduous beta diversity approaches; while revealing the taxonomic identities of the communities they structure (Hernández Limón et al. *in review*). The dominant axis of cross-block microbial organization in the Laurentian Great Lakes is thermal stratification, which partitions warm and cold water assemblages that are largely non-overlapping and taxonomically irreplaceable. The organisms that define cold water habitats — chemolithotrophic specialists, silica-dependent diatoms, cold-adapted flagellates — have no warm water equivalents, while warm surface communities dominated by streamlined oligotrophs, copiotrophic heterotrophs, and mixotrophic flagellates are poised to expand as stratification intensifies (O’Reilly et al., 2015; Woolway et al., 2021). These findings suggest that warming-driven changes in stratification will restructure the Great Lakes microbial system in taxonomically specific and potentially irreversible ways. The LDA-GCA framework developed here provides a replicable template for characterizing microbial community change across large lake systems globally, where rapid warming and stratification intensification are restructuring aquatic ecosystems (O’Reilly et al., 2015; Woolway et al., 2021).

## Data and Methods

### Sample collection, DNA Processing, and Metadata

Water samples were collected aboard the R/V Lake Guardian from 2012–2019 across the five Laurentian Great Lakes during biannual spring and summer cruises at 15–17 stations and 3–4 depths per station (Supp. Fig. 1). Sample collection, DNA extraction, sequencing, and metadata compilation are described in detail in Hernández Limón et al. *in review*. Briefly, water was sequentially filtered through 1.6 μm (Whatman GF/A) and 0.2 μm (Millipore Express Plus) filters, yielding particle-associated (PA) and free-living (FL) prokaryotic fractions, and large (LSF) and small (SSF) chloroplast-containing eukaryotic fractions. At Lake Erie Spring stations, an additional 5 μm filter (Durapore PVDF) was collected; both prokaryotic and chloroplast communities on this filter were excluded from analyses to maintain consistent size fraction definitions across all lakes.

Raw sequences were processed in QIIME2 (v2026.1; Bolyen et al., 2019) using DADA2 (Callahan et al., 2016) to generate amplicon sequence variants (ASVs). 16S and 18S reads were separated and classified against custom Silva 138.2 (Quast et al., 2013) and PR2 (v5; Guillou et al., 2013) databases following the primer-specific pipeline described in McNichol et al. (2021) and Yeh et al. (2021). Prokaryotic ASVs (Bacteria and Archaea, excluding mitochondria and chloroplasts) and chloroplast ASVs (photosynthetic eukaryotes) were analyzed separately. Taxonomy was assigned using Silva138.2 at a 0.70 confidence threshold; chloroplast ASVs were further annotated using PR2. ASVs with fewer than 10 reads across all samples, those detected in only a single sample, and samples with fewer than 1,000 reads were removed. Environmental metadata were provided by the EPA and supplemented with a 14-day mean air temperature from the National Solar Radiation Database and estimated light penetration from surface Global Horizontal Irradiance; full derivation is described in Hernández Limón et al. *in review*.

All analyses were conducted in R (v4.5.3). Visualizations were produced using ggplot2 (v3.5.1), cowplot (v1.1.3), and ggpubr (v0.6.0).

### Subcommunity identification with *alto*

Subcommunities were identified using Latent Dirichlet Allocation (LDA) implemented in the R package alto (v0.1.0), which includes diagnostic algorithms for selecting the optimal number of topics (k) (Fukuyama et al., 2023; Sankaran & Holmes, 2019). ASV counts were modeled across k = 1–25, with subcommunities aligned using both the product and transport methods. Optimal k was selected based on leveling off of path counts and refinement scores combined with declining coherence at higher k (Supp. Fig. 2) and validated using perplexity scores in topicmodels (v0.2-16) with a two-thirds training and one-third validation split. Final k values were FL = 10, PA = 10, SSF = 9, LSF = 11. LDA produced two outputs used in subsequent analyses: subcommunity distributions across samples (γ) and taxon distributions across subcommunities (β).

### Subcommunity Characteristics

Shannon entropy was calculated for each sample from γ using the entropy package (v1.3.1; Hausser & Strimmer, 2009) to assess the evenness of subcommunity contributions within samples. Higher entropy reflects more even mixing across subcommunities; lower entropy indicates dominance by few subcommunities. Differences in entropy distributions between biological blocks were assessed using two-sample Kolmogorov-Smirnov tests, with p < 0.001 considered significant. Subcommunity distributions were visualized as stacked bar plots of γ proportions with samples hierarchically clustered using Ward’s method (Supp. Fig 3).

### Fingerprint Taxa Identification with GoM-DE

Fingerprint taxa for each subcommunity were identified using Grade of Membership Differential Expression analysis (GoM-DE) in the fastTopics package (Carbonetto et al., 2023), applied independently to each fraction. GoM-DE used γ from the alto LDA as weights, fitting Poisson regressions for each ASV to estimate topic-specific enrichment relative to all other topics, with raw ASV counts as input and a pseudocount of 0.1. Effect sizes are reported as log2 fold-change (log2FC) using the “le” statistic with local false sign rate (lfsr) shrinkage. ASVs with lfsr < 0.05 and log2FC > 0 were retained as significantly enriched, and the top 30 per topic selected as fingerprint taxa. Topic specificity was assessed by tracking the number of topics in which each ASV was significantly enriched. Visualization focused on the top 25 fingerprint ASVs exclusive to a single topic as the most diagnostic markers of subcommunity identity.

### Environmental Associations with Limma

Significant associations between subcommunity proportions and environmental variables were identified using limma (Ritchie et al., 2015) applied independently to each fraction. Subcommunity proportions (γ) were centered log-ratio (CLR) transformed jointly across all topics per sample prior to modeling, with a pseudocount of 1×10⁻⁶. CLR-transformed proportions were modeled against 12 scaled continuous environmental variables simultaneously using a single design matrix, with empirical Bayes shrinkage via eBayes applied to stabilize variance estimates across topics. Only samples with complete environmental metadata were retained. Effect sizes are reported as CLR-space coefficients reflecting magnitude and direction of association. Associations were considered significant at BH-adjusted p < 0.05, with correction applied within each environmental variable across topics.

### Environmental Prediction with Random Forest and SHAP

Random forest (RF) models were fit using the ranger package (v0.16.0; Wright & Ziegler, 2017) separately for each topic within each fraction, with CLR-transformed γ as the outcome and 12 scaled environmental predictors as inputs. Models used 500 trees, mtry set to the square root of the number of predictors, and minimum node size of 5. Model performance was assessed using out-of-bag R² (OOB R²). Variable importance and direction were assessed using SHAP values computed with fastshap (v0.1.1; Greenwell, 2023) with 500 simulations per model. Mean absolute SHAP values were scaled 0–1 within each topic to facilitate visual comparison. Direction was determined by Spearman correlation between feature values and SHAP values, with |ρ| > 0.1 used as a visualization threshold rather than a formal significance criterion.

### Cross-fraction Integration with Generalized Canonical Analysis

Generalized Canonical Analysis (GCA) was applied to identify shared axes of variation across five blocks: free-living prokaryotes (FL), particle-associated prokaryotes (PA), small chloroplast-containing eukaryotes (SSF), large chloroplast-containing eukaryotes (LSF), and environmental variables (Env). GCA identifies linear combinations of variables within each block that maximize cross-block covariance relative to within-block covariance, enabling direct comparison across datasets that cannot otherwise be jointly analyzed. Topic proportions (γ) from each microbial block were ILR-transformed using the compositions package (v2.0-9; van den Boogaart & Tolosana-Delgado, 2013); environmental variables were scaled to zero mean and unit variance with missing values replaced by zero.

All blocks were aligned to a common set of 678 samples using FL as the reference. Samples in FL but absent from PA were excluded. For SSF and LSF, samples lacking detectable eukaryotic sequences were retained and assigned a placeholder topic (SSF_0 or LSF_0, proportion = 1) encoding biological absence as an explicit compositional state rather than missing data, as absence of chloroplast-containing eukaryotes reflects genuine ecological conditions including cold isothermal and deep low-light environments. Of the 678 samples, 409 lacked SSF detection and 120 lacked LSF detection, consistent with the restriction of small eukaryotes to thermally stratified productive conditions in the Great Lakes. GCA was implemented by solving a generalized eigenvalue problem comparing the global covariance matrix (Σ) to a block-diagonal within-block covariance matrix (Σ₀), regularized with a ridge penalty (λ = 10). Five canonical axes were retained based on the scree plot of proportional variance explained, corresponding to the number of input blocks. Block contributions were summarized as mean absolute canonical variates per block per axis, and environmental loadings used to interpret each axis ecologically.

### Subcommunity Composition Across Space and Time

Subcommunity composition across samples was visualized using stacked tile plots adapted from Symul et al. (2023), in which each sample (ordered by station and year) is shown on the x-axis and each subcommunity on the y-axis, with tile height and size scaled proportionally to the subcommunity’s contribution to that sample. Only subcommunities contributing more than their proportional share (threshold: 3/k, where k is the number of subcommunities) are displayed, highlighting dominant assemblages while suppressing background contributions. Samples are grouped by station and ordered west to east across the basin, with years 2012–2019 nested within each station, allowing simultaneous visualization of spatial, temporal, and subcommunity structure across the dataset.

## Data Availability

Sequences are deposited in NCBI under BioProject accession number PRJNA1259575 and will be made publicly available upon peer-reviewed publication of the companion manuscript. A toy dataset and fully documented analysis code reproducing all manuscript figures are available at https://github.com/MDHDZ91/greatlakes-lda-microbial-communities.

## Acknowledgements

This work was supported by Illinois–Indiana Sea Grant (NA14OAR4170095, subaward 2014-02342-05) and the National Science Foundation Biological Oceanography Program (OCE-1830011) to M.L.C. M.D.H.L. was supported by the Neubauer Family Doctoral Fellowship (2018–2019 cohort), the NSF National Research Traineeship program (DGE-1735359), the Ford Foundation Predoctoral Fellowship (2020 Award), and the NITMB Traineeship Program (NSF DMS-2235451; Simons Foundation MPS-NITMB-00005320). C.D acknowledges support by the National Science Foundation under Award Numbers IIS:2238616, and IOS:2520677. F.B acknowledges support by the National Science Foundation under Award Number IOS:2520677. Computational analyses were performed using resources provided by the University of Chicago Research Computing Center.

## Supplementary figures

**Supp Fig 1.**
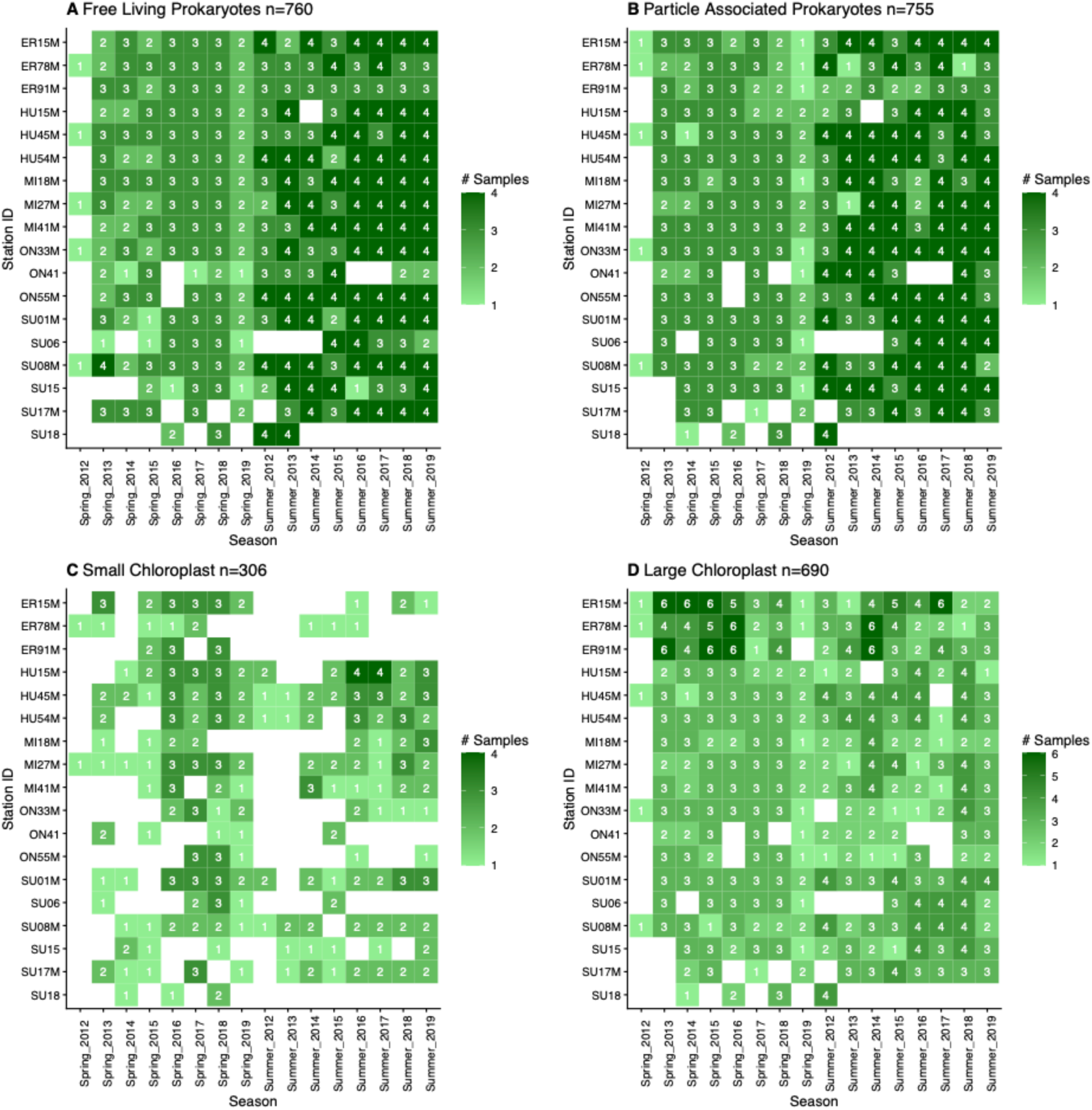
Sampling effort for LDA analysis across years (x-axis) and stations (y-axis) for each dataset: (A) Free Living Prokaryotes, (B) Particle Associated Prokaryotes, (C) Small size fraction chloroplasts, and (D) Large size fraction chloroplasts. Color indicates the number of samples per station-year.

**Supp Fig. 2.**
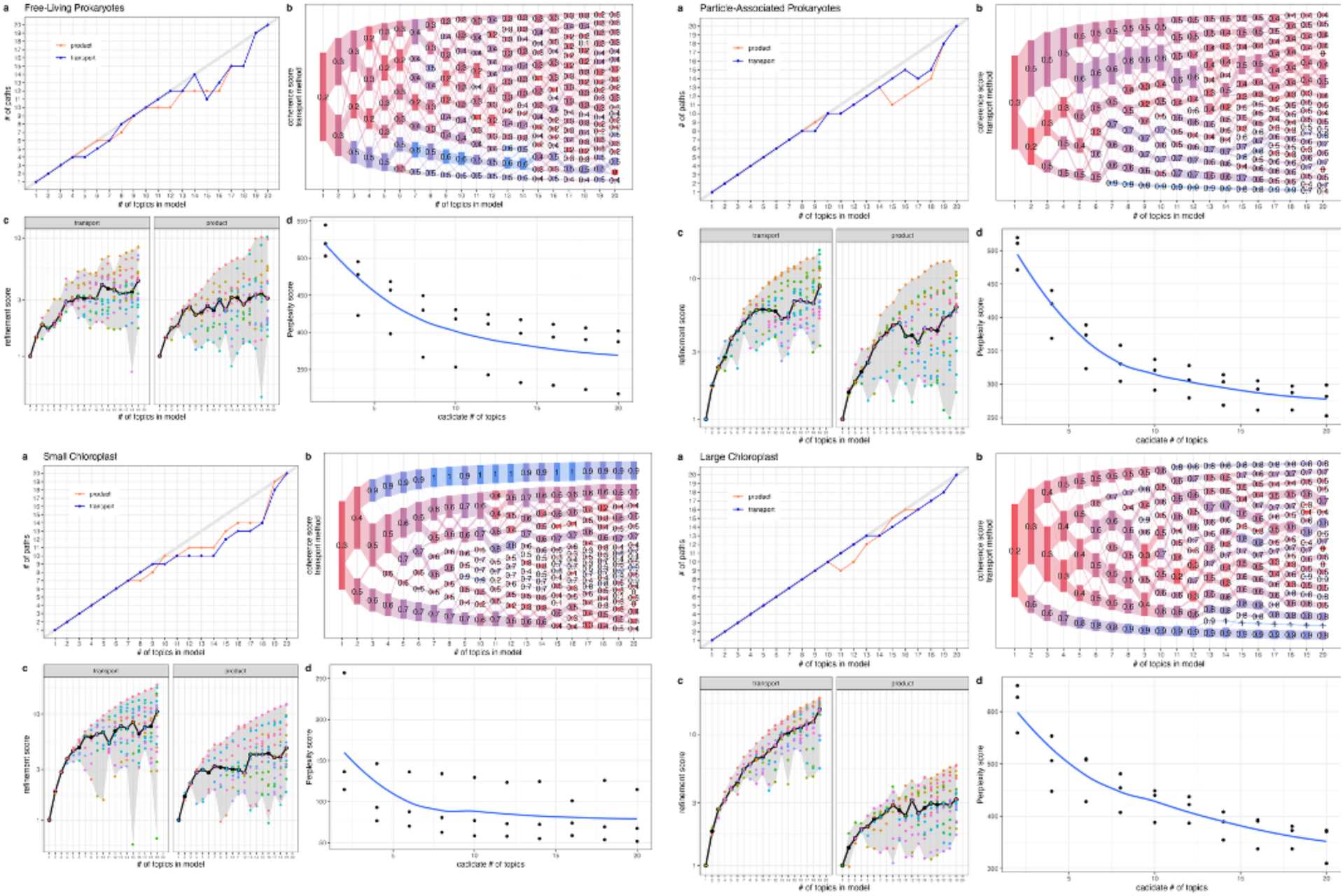
LDA model diagnostics used to select the number of topics (subcommunities, SCs) for each size fraction. Each panel shows four diagnostic plots: (top left) number of paths, with lines indicating performance from product- and transport-based methods; (bottom left) refinement score for each method; (top right) coherence score; and (bottom right) perplexity.

**Supp Fig. 3.**
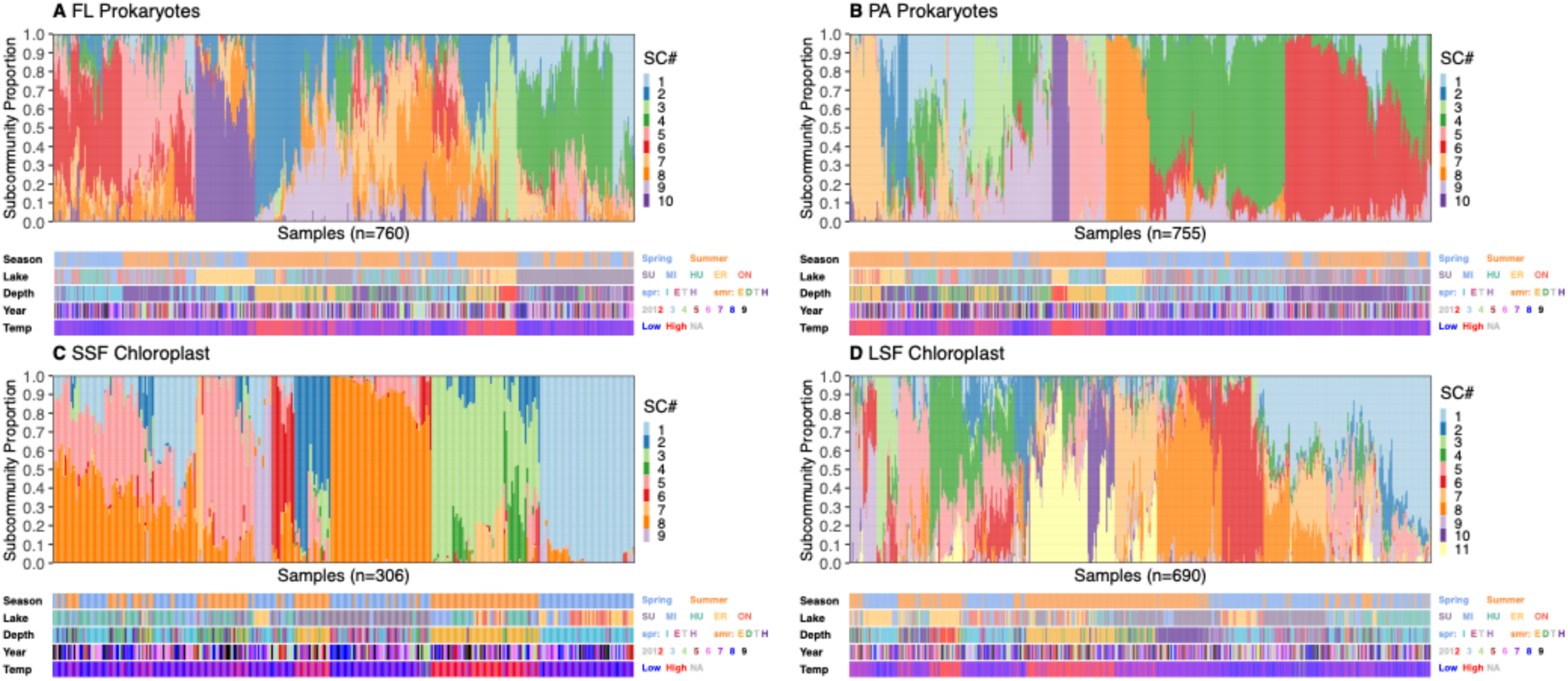
Sample-by-subcommunity (SC) composition for different blocks with samples ordered by Ward clustering and colored by SC identity. Metadata tracks below show season, lake, depth, year, and temperature.

**Supp Fig. 4.**
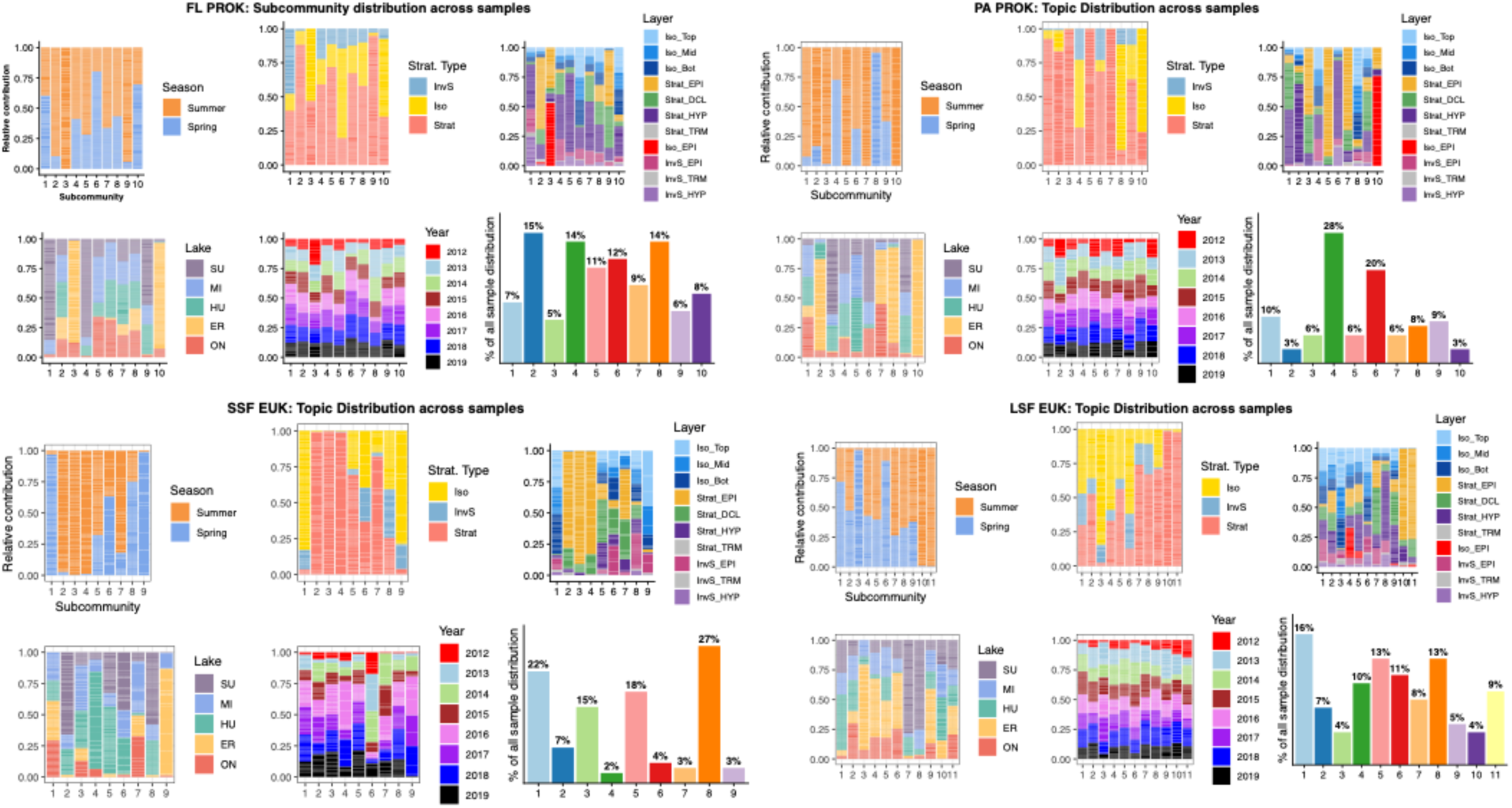
Environmental and spatial context of microbial subcommunities. Panels show the relative contribution of each SC across samples, grouped by metadata categories including season, stratification type, water column strata, lake, and year. The final bar (far right) shows the percentage of all samples in which each SC was detected, indicating sub-community prevalence across the dataset.

**Supp. Fig 5.**
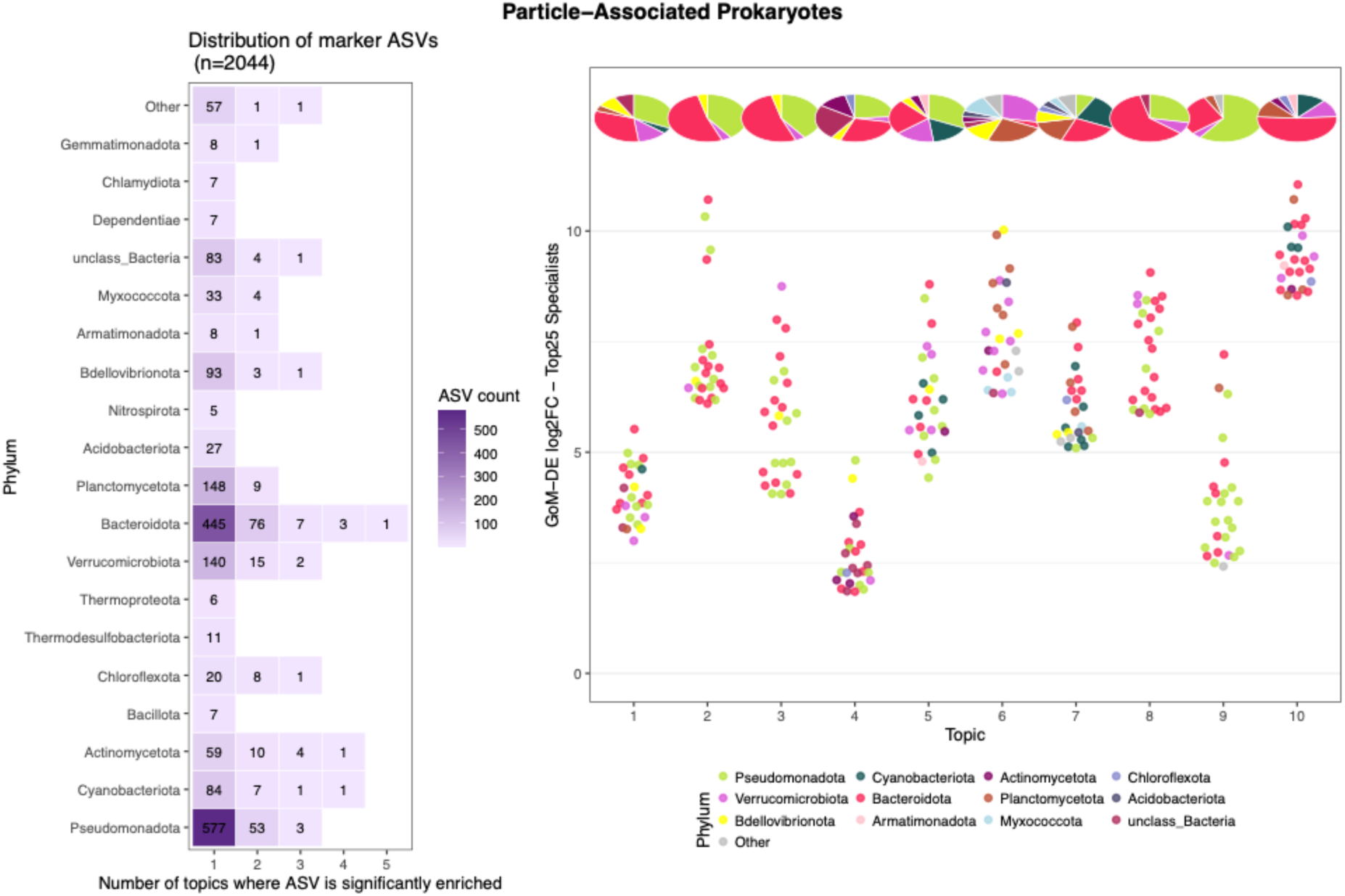
GoMDE distributions. (left) Distribution of all marker ASVs (n=2,044) by the number of SCs in which each ASV was significantly enriched, (right) log2 fold-change of the top 25 specialist ASVs per SC. Points are colored by phylum; pie charts above each SC show the phylum composition of all specialist ASVs for that topic.

**Supp. Fig 6.**
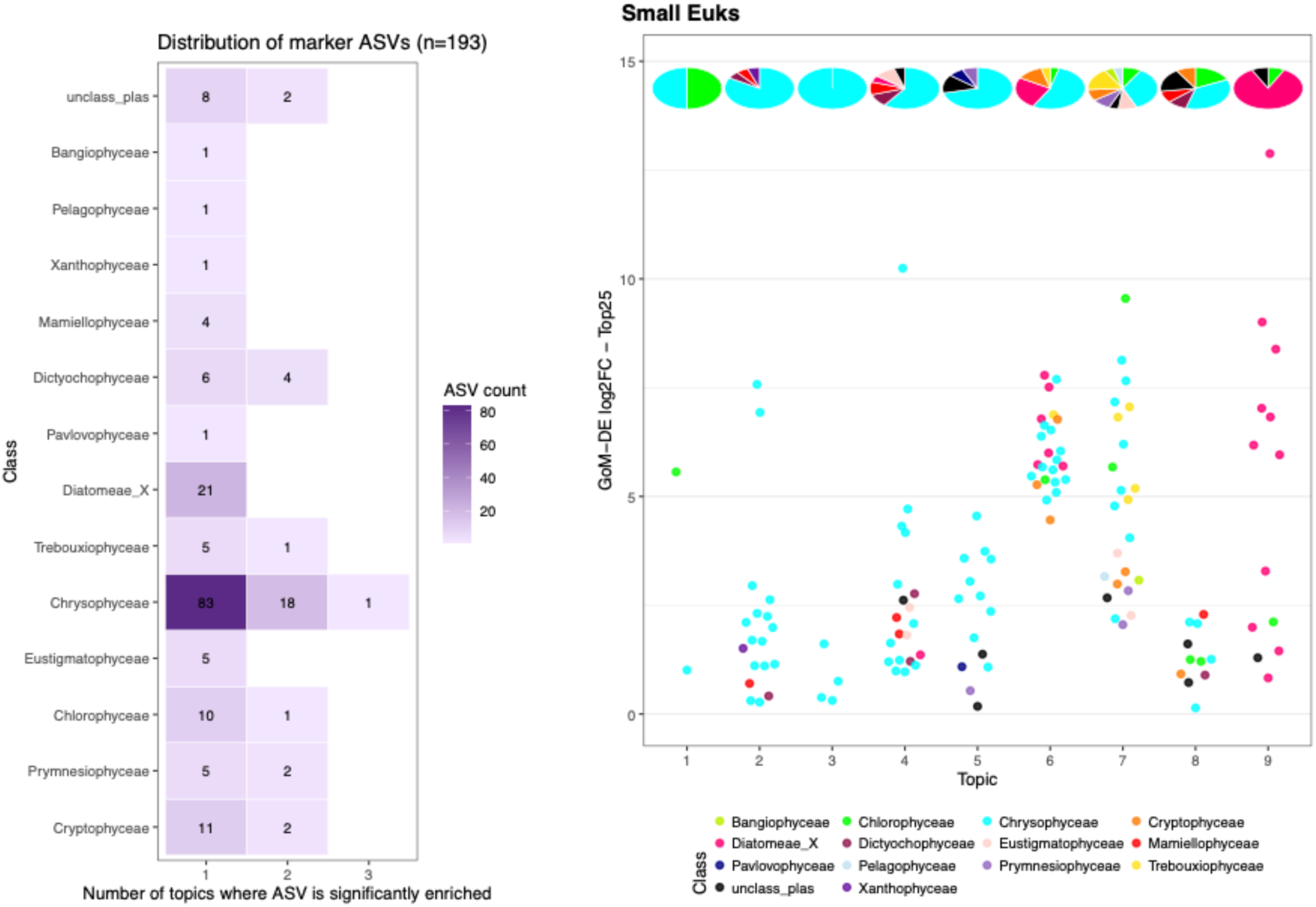
GoMDE distributions. (left) Distribution of all marker ASVs (n=2,044) by the number of SCs in which each ASV was significantly enriched, (right) log2 fold-change of the top 25 specialist ASVs per SC. Points are colored by phylum; pie charts above each SC show the phylum composition of all specialist ASVs for that topic.

**Supp. Fig 7.**
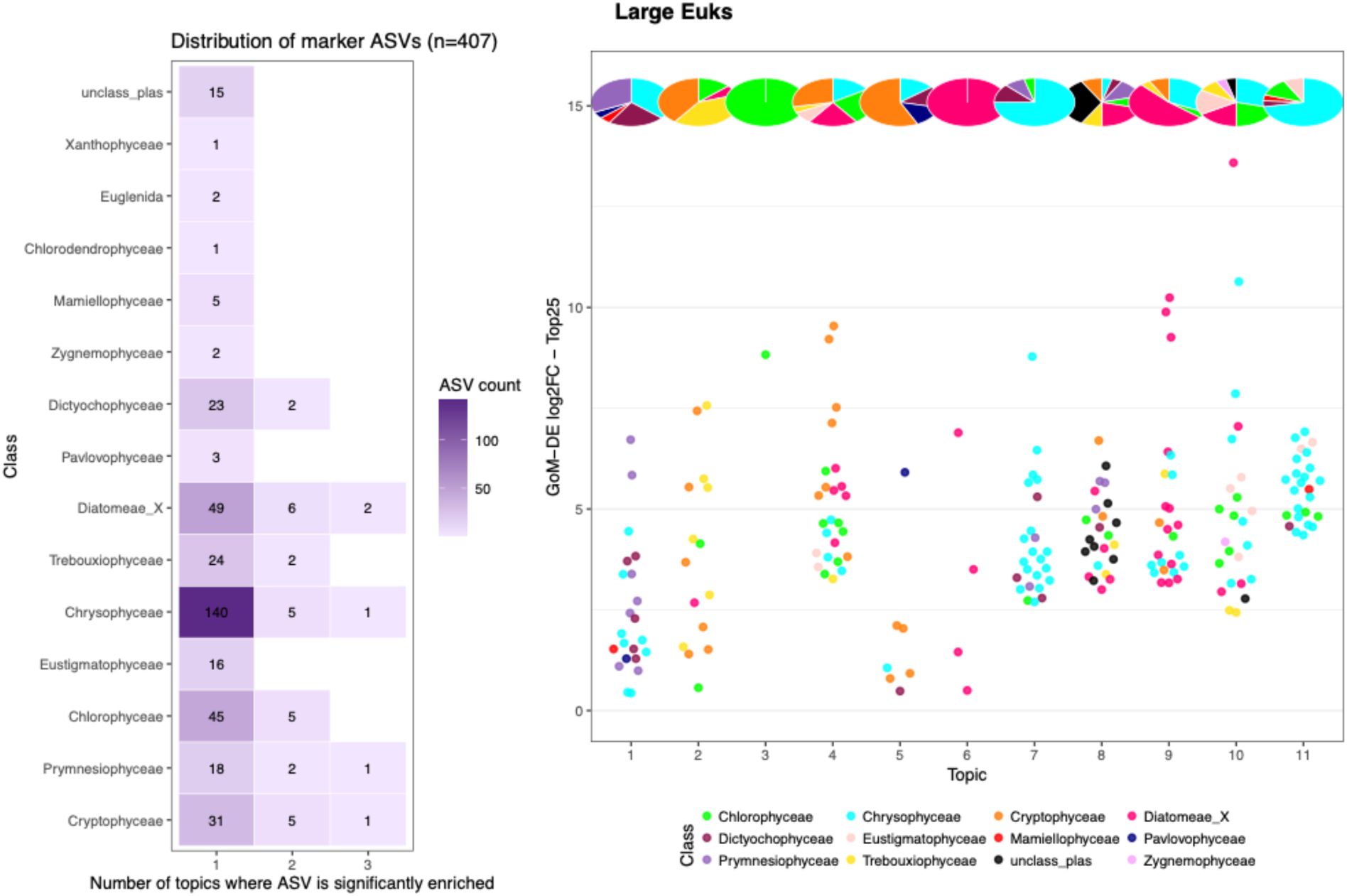
GoMDE distributions. (left) Distribution of all marker ASVs (n=2,044) by the number of SCs in which each ASV was significantly enriched, (right) log2 fold-change of the top 25 specialist ASVs per SC. Points are colored by phylum; pie charts above each SC show the phylum composition of all specialist ASVs for that topic.

**Supp. Fig. 8.**
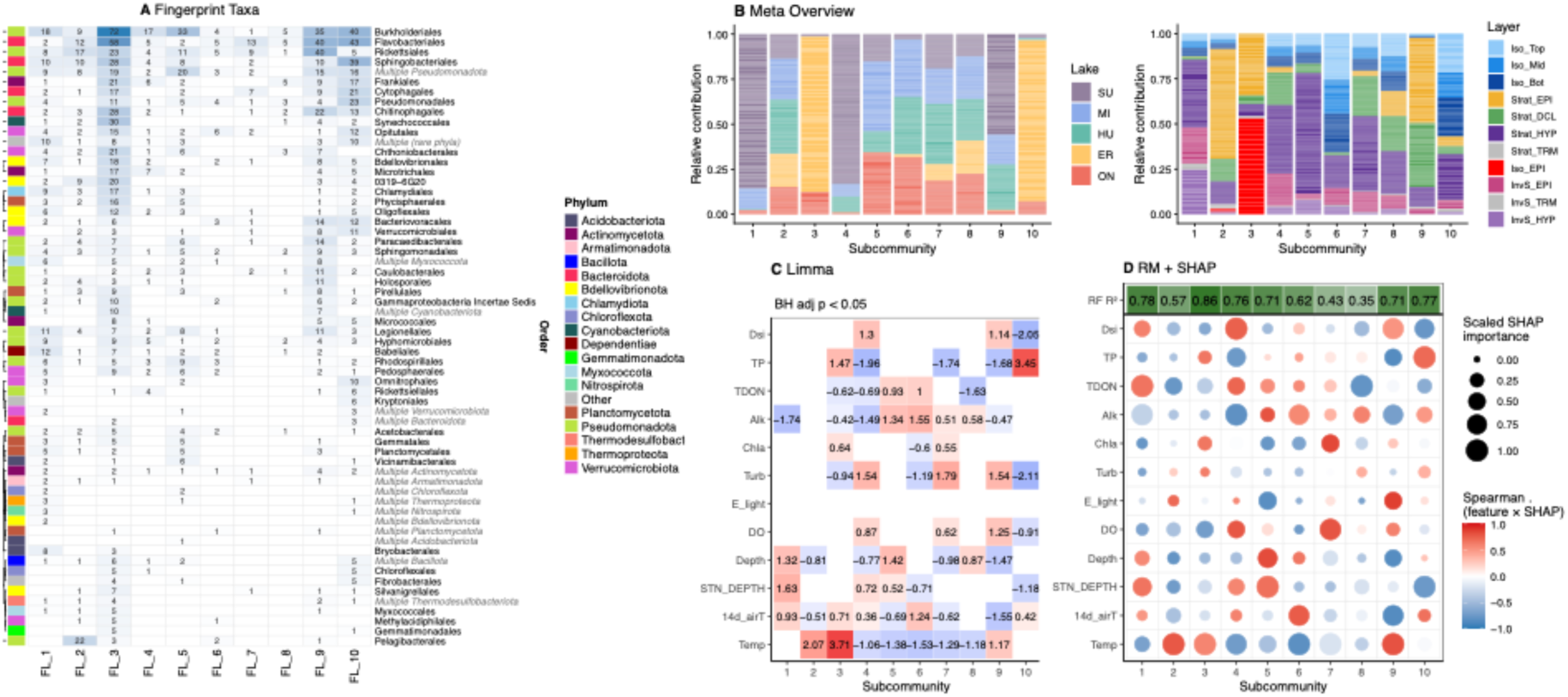
Taxonomic identity and environmental associations of free-living prokaryote (FL) subcommunities. (A) Fingerprint taxa identified by GoM-DE discriminant analysis. Each column represents one subcommunity (SC) and each row represents a bacterial order. Numbers indicate the count of differentially enriched ASVs (BH-adjusted p < 0.05) from that order in that SC, colored by phylum. SC-specific ASVs (unique to one SC) are shown in bold; orders appearing in italic represent multi-phylum groupings. (B) Metadata overview showing the relative contribution of each SC across samples, summarized by lake identity (left) and water column layer (right). (C) Limma linear model results showing significant associations (BH-adjusted p < 0.05) between SC proportions and environmental variables. Numbers indicate standardized effect sizes; red indicates positive association and blue indicates negative association. (D) Random forest predictive importance (RF R^2^ shown in header) and SHAP-based environmental driver analysis. Circle size reflects scaled SHAP importance and color reflects the Spearman correlation between feature value and SHAP value, indicating direction of effect.

**Supp. Fig. 9.**
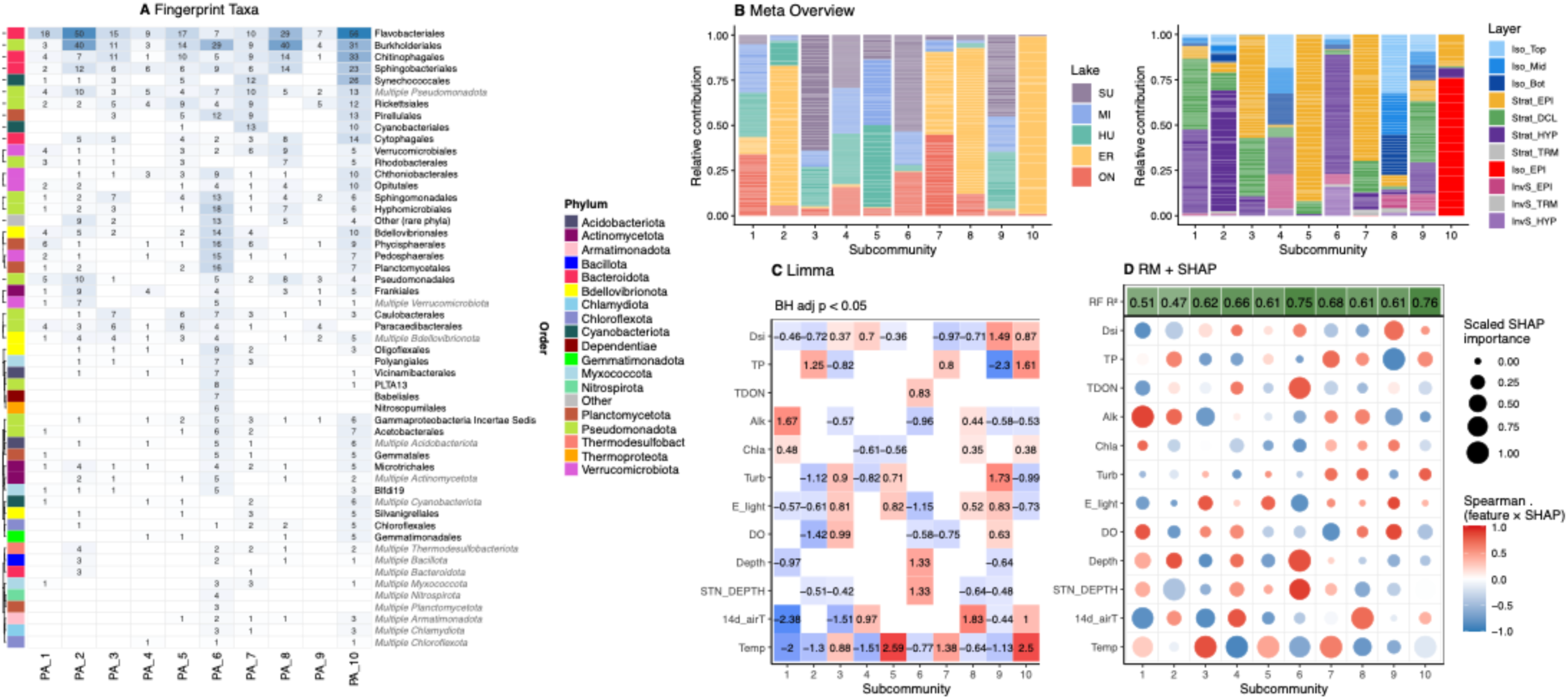
Taxonomic identity and environmental associations of particle-associated prokaryote (PA) subcommunities. (A) Fingerprint taxa identified by GoM-DE discriminant analysis. Each column represents one subcommunity (SC) and each row represents a bacterial order. Numbers indicate the count of differentially enriched ASVs (BH-adjusted p < 0.05) from that order in that SC, colored by phylum. SC-specific ASVs (unique to one SC) are shown in bold; orders appearing in italic represent multi-phylum groupings. (B) Metadata overview showing the relative contribution of each SC across samples, summarized by lake identity (left) and water column layer (right). (C) Limma linear model results showing significant associations (BH-adjusted p < 0.05) between SC proportions and environmental variables. Numbers indicate standardized effect sizes; red indicates positive association and blue indicates negative association. (D) Random forest predictive importance (RF R^2^ shown in header) and SHAP-based environmental driver analysis. Circle size reflects scaled SHAP importance and color reflects the Spearman correlation between feature value and SHAP value, indicating direction of effect.

**Supp. Fig. 10.**
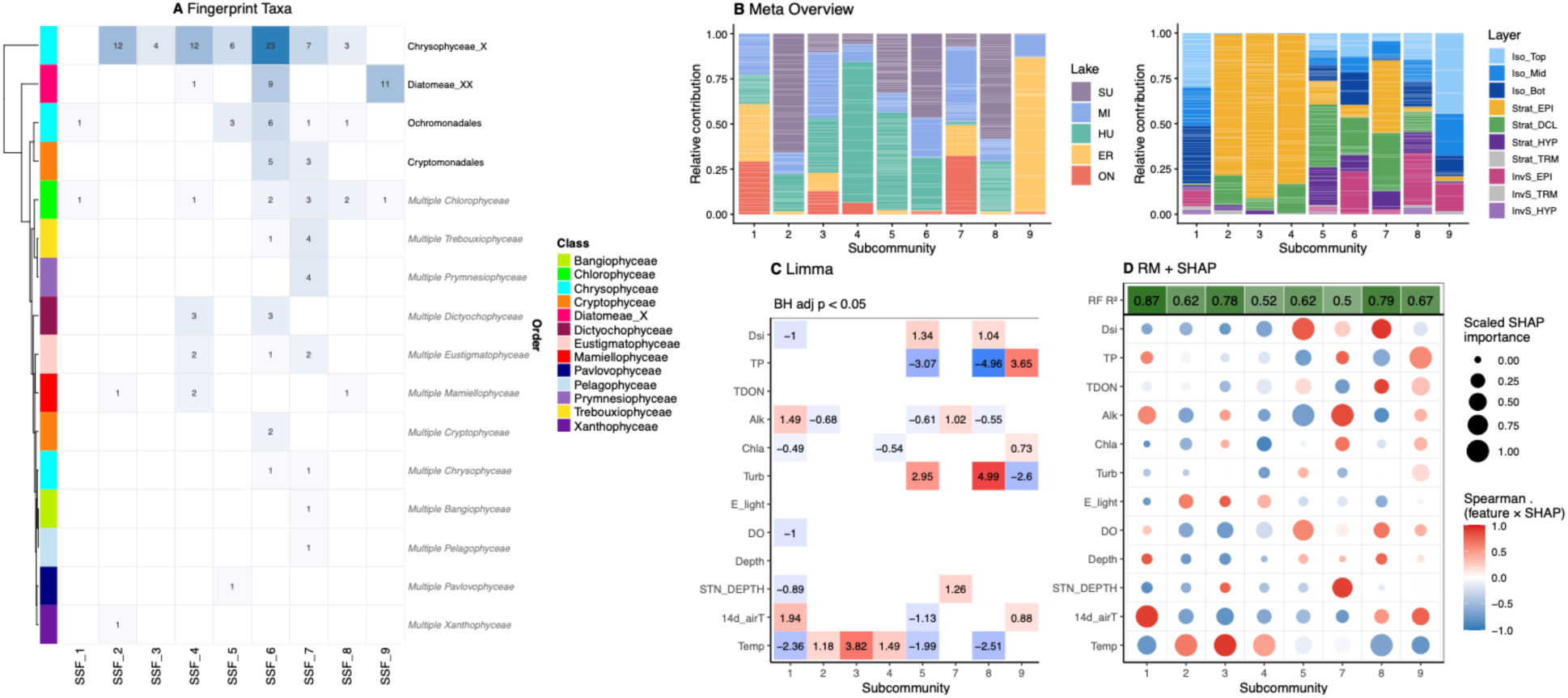
Taxonomic identity and environmental associations of small size fraction chloroplast (SSF) subcommunities. (A) Fingerprint taxa identified by GoM-DE discriminant analysis. Each column represents one subcommunity (SC) and each row represents a phytoplankton order. Numbers indicate the count of differentially enriched ASVs (BH-adjusted p < 0.05) from that class in that SC, colored by taxonomic class. SC-specific ASVs (unique to one SC) are shown in bold; classes appearing in italic represent multi-class groupings. (B) Metadata overview showing the relative contribution of each SC across samples, summarized by lake identity (left) and water column layer (right). (C) Limma linear model results showing significant associations (BH-adjusted p < 0.05) between SC proportions and environmental variables; SSF 6 is excluded as no associations passed the significance threshold. Numbers indicate standardized effect sizes; red indicates positive association and blue indicates negative association. (D) Random forest predictive importance (RF R^2^ shown in header) and SHAP-based environmental driver analysis; SSF 6 is excluded (RF R^2^ = 0.30, the lowest predictability observed across all SCs in the study), suggesting this subcommunity is not strongly structured by the measured environmental variables. Circle size reflects scaled SHAP importance and color reflects the Spearman correlation between feature value and SHAP value, indicating direction of effect.

**Supp. Fig. 11.**
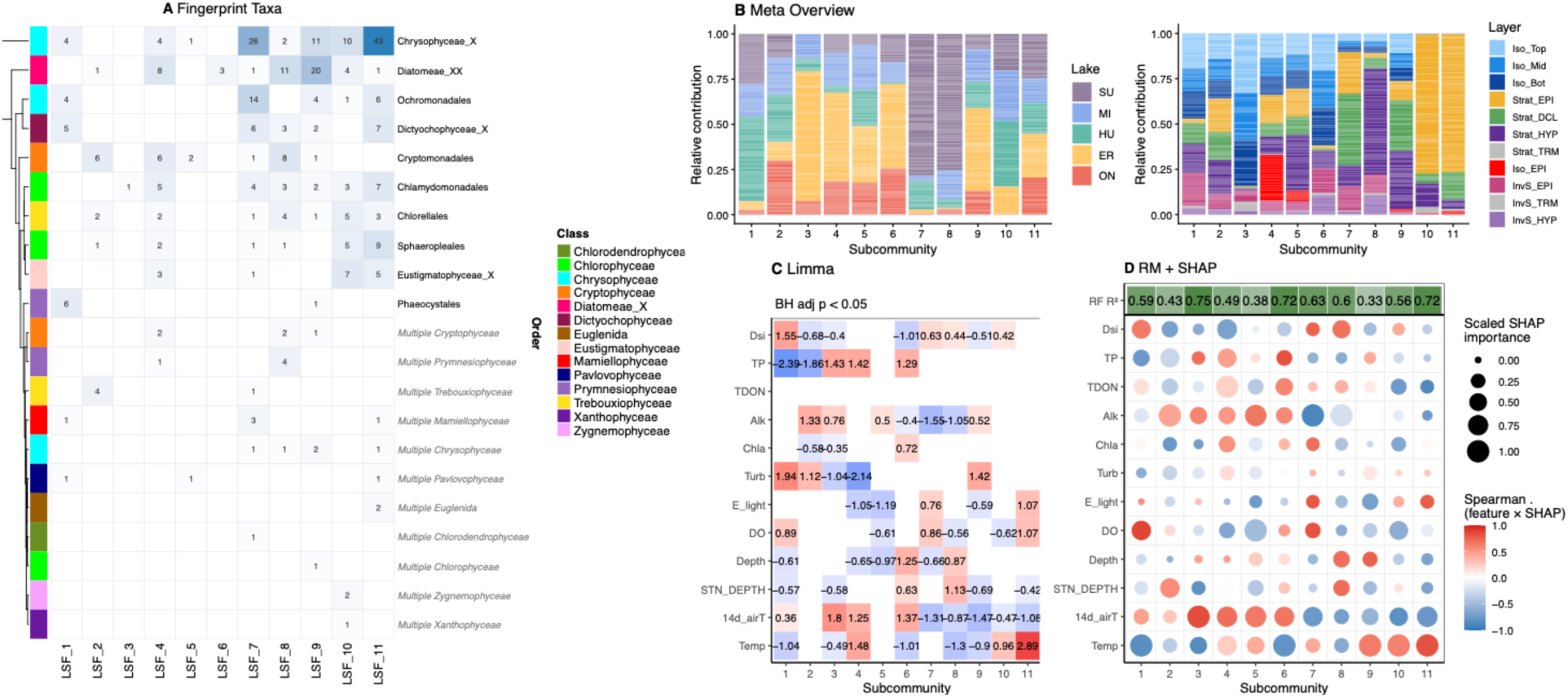
Taxonomic identity and environmental associations of large size fraction chloroplast (LSF) subcommunities. (A) Fingerprint taxa identified by GoM-DE discriminant analysis. Each column represents one subcommunity (SC) and each row represents a phytoplankton order. Numbers indicate the count of differentially enriched ASVs (BH-adjusted p < 0.05) from that order in that SC, colored by class. SC-specific ASVs (unique to one SC) are shown in bold; orders appearing in italic represent multi-class groupings. (B) Metadata overview showing the relative contribution of each SC across samples, summarized by lake identity (left) and water column layer (right). (C) Limma linear model results showing significant associations (BH-adjusted p < 0.05) between SC proportions and environmental variables. Numbers indicate standardized effect sizes; red indicates positive association and blue indicates negative association. (D) Random forest predictive importance (RF R^2^ shown in header) and SHAP-based environmental driver analysis. Circle size reflects scaled SHAP importance and color reflects the Spearman correlation between feature value and SHAP value, indicating direction of effect.

**Supp. Fig. 12.**
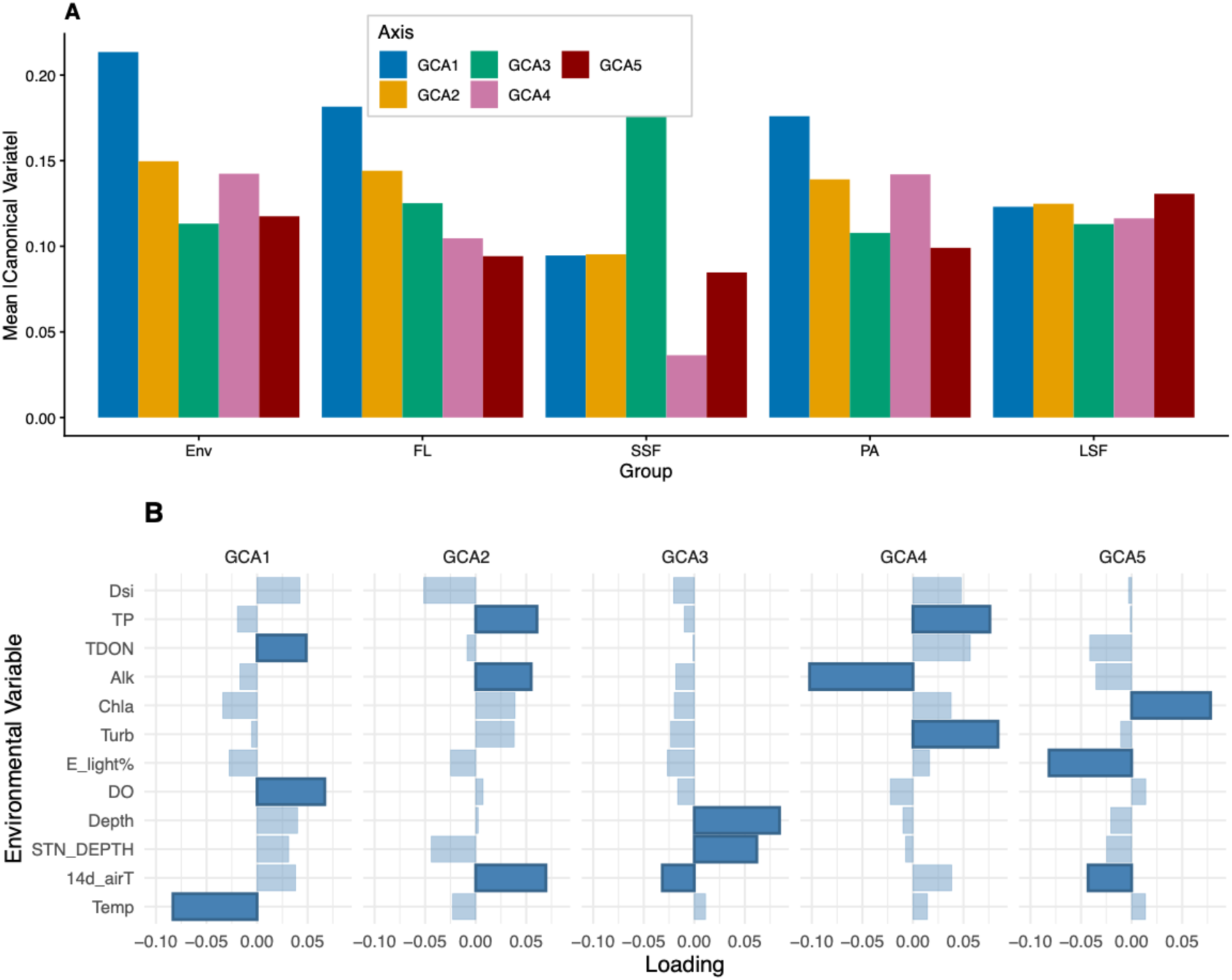
Group Compositional Analysis (GCA) of subcommunity distributions across all four biological blocks. (A) Mean absolute canonical variate scores for each GCA axis (GCA1-GCA5) across the five variable groups — environmental variables (Env), free- living prokaryotes (FL), small size fraction chloroplasts (SSF), particle-associated prokaryotes (PA), and large size fraction chloroplasts (LSF) — reflecting the relative contribution of each group to each axis. (B) Environmental variable loadings for each GCA axis. Bar length reflects the magnitude of each variable’s loading and direction indicates positive or negative association.

## Notes

### Competing Interest Statement

The authors have declared no competing interest.

### Summary of Updates

The acknowledgments section was missing funding sources as well as their award numbers. We have added that information for all authors.

